# Structural and functional characterization of the Spo11 core complex

**DOI:** 10.1101/2020.02.21.960211

**Authors:** Corentin Claeys Bouuaert, Sam E. Tischfield, Stephen Pu, Eleni P. Mimitou, Ernesto Arias-Palomo, James M. Berger, Scott Keeney

**Author notes:** Correspondence (C.C.B.), (S.K.). ST: Human Oncology and Pathology Program, Memorial Sloan Kettering Cancer Center, New York, US. EPM: New York Genome Center, New York, US. EAP: Department of Structural & Chemical Biology, CIB Margarita Salas (CSIC) 28040 Madrid, Spain. CCB: Louvain Institute of Biomolecular Science and Technology, Université catholique de Louvain, Louvain-La-Neuve, Belgium.

## Abstract

Spo11, which makes DNA double-strand breaks (DSBs) essential for meiotic recombination, is poorly understood mechanistically because it has been recalcitrant to biochemical study. Here, we provide a molecular analysis of *S. cerevisiae* Spo11 purified with partners Rec102, Rec104 and Ski8. Rec102 and Rec104 jointly resemble the B subunit of archaeal Topoisomerase VI, with Rec104 similar to a GHKL domain but without conserved ATPase motifs. Unexpectedly, the Spo11 complex is monomeric (1:1:1:1 stoichiometry), indicating that dimerization may control DSB formation. Reconstitution of DNA binding reveals topoisomerase-like preferences for duplex-duplex junctions and bent DNA. Spo11 also binds noncovalently but with high affinity to DNA ends mimicking cleavage products, suggesting a mechanism to cap DSB ends. Mutations that reduce DNA binding *in vitro* attenuate DSB formation, alter DSB processing, and reshape the DSB landscape *in vivo*. Our data reveal structural and functional similarities between the Spo11 core complex and Topo VI, but also highlight differences reflecting their distinct biological roles.

## Introduction

Meiosis in most sexually reproducing organisms requires recombination that is initiated when Spo11 introduces DNA double-strand breaks (DSBs) (Lam and Keeney, 2015). Spo11 evolved from the DNA cleaving subunit of an archaeal type IIB topoisomerase (Bergerat et al., 1997; Keeney et al., 1997). Topo VI, the archetypal member of this family, is a tetramer of two A subunits (DNA cleaving) and two B subunits (GHKL-type ATPase) (Corbett et al., 2007; Graille et al., 2008). DNA breakage involves concerted action of the A subunits attacking opposite strands of a duplex using active-site tyrosines to produce covalent 5’-phosphotyrosyl links (Buhler et al., 2001). Topo VI coordinates the transient formation of a DSB with passage of an intact duplex through the break in order to modulate DNA topology.

In contrast, Spo11 remains trapped in the covalent complex, producing the breaks that initiate recombination (Lam and Keeney, 2015). Endonucleolytic cleavage of each Spo11-bound strand releases Spo11 along with a short oligonucleotide still covalently attached to the active-site tyrosine (Spo11-oligo complexes) (Neale et al., 2005). This cleavage frees DSB ends for further exonucleolytic resection followed by homology search and strand exchange (Symington, 2014; Hunter, 2015).

Spo11 requires a cohort of partners (Lam and Keeney, 2015). Their sequences are highly diverged (Keeney, 2008), but studies in several species showed that most are in fact conserved functionally (Kumar et al., 2010; Miyoshi et al., 2012; Robert et al., 2016; Stanzione et al., 2016; Vrielynck et al., 2016; Tesse et al., 2017). One important development was the discovery in mouse and plants of a Spo11 partner homologous to the Topo VI B subunit (Top6BL) (Robert et al., 2016; Vrielynck et al., 2016). This suggested that the meiotic DSB machinery is more similar to the ancestral topoisomerase than previously thought. Discovery of Top6BL further led to identification of putative B-type subunits in other organisms, including yeasts and flies. This protein in *S. cerevisiae*, Rec102, was long known to be important for Spo11 activity (Malone et al., 1991; Kee and Keeney, 2002; Jiao et al., 2003; Maleki et al., 2007), but its potential relationship with the B-type subunit had remained unnoticed. Intriguingly, structural modeling suggested that the B-type subunits of *S. cerevisiae, S. pombe* and *Drosophila* lack the GHKL domain that mediates ATP-dependent dimerization in Topo VI, although a degenerate version of this domain is found in the mouse and plant counterparts (Robert et al., 2016; Vrielynck et al., 2016).

Despite >20 years of progress since the discovery of Spo11 function (Bergerat et al., 1997; Keeney et al., 1997), biochemical studies of the meiotic DSB machinery have been lacking, largely due to difficulties in isolating key components. In the absence of biochemical and structural information, how Spo11 engages DNA, its functional relationships with Topo VI, and the roles of essential Spo11-accessory factors have remained a matter of conjecture. Here, we present the first *in vitro* characterization of a stoichiometric, Spo11-containing core complex from yeast. We elucidate similarities of the structure and DNA-binding properties of the complex with the ancestral Topo VI but also uncover unique features that are presumably related to its exaptation as the meiotic DSB generator.

## Results

### The meiotic DSB core complex has a 1:1:1:1 stoichiometry

Yeast Spo11 most closely associates with Ski8, Rec102, and Rec104 (Kee and Keeney, 2002; Jiao et al., 2003; Arora et al., 2004; Kee et al., 2004; Prieler et al., 2005; Sasanuma et al., 2007; Lam and Keeney, 2015), so we co-expressed these four proteins in insect cells and purified soluble complexes (**Figure 1A**). We used a C-terminal tag on Spo11 that is functional *in vivo (Kugou et al., 2009)* and purified complexes with or without maltose binding protein (MBP) fused at the N-terminus of Rec102. Omission of any of the other proteins resulted in low Spo11 solubility (**Supplementary Figure 1**). We refer to this as the meiotic DSB core complex.

**Figure 1:**
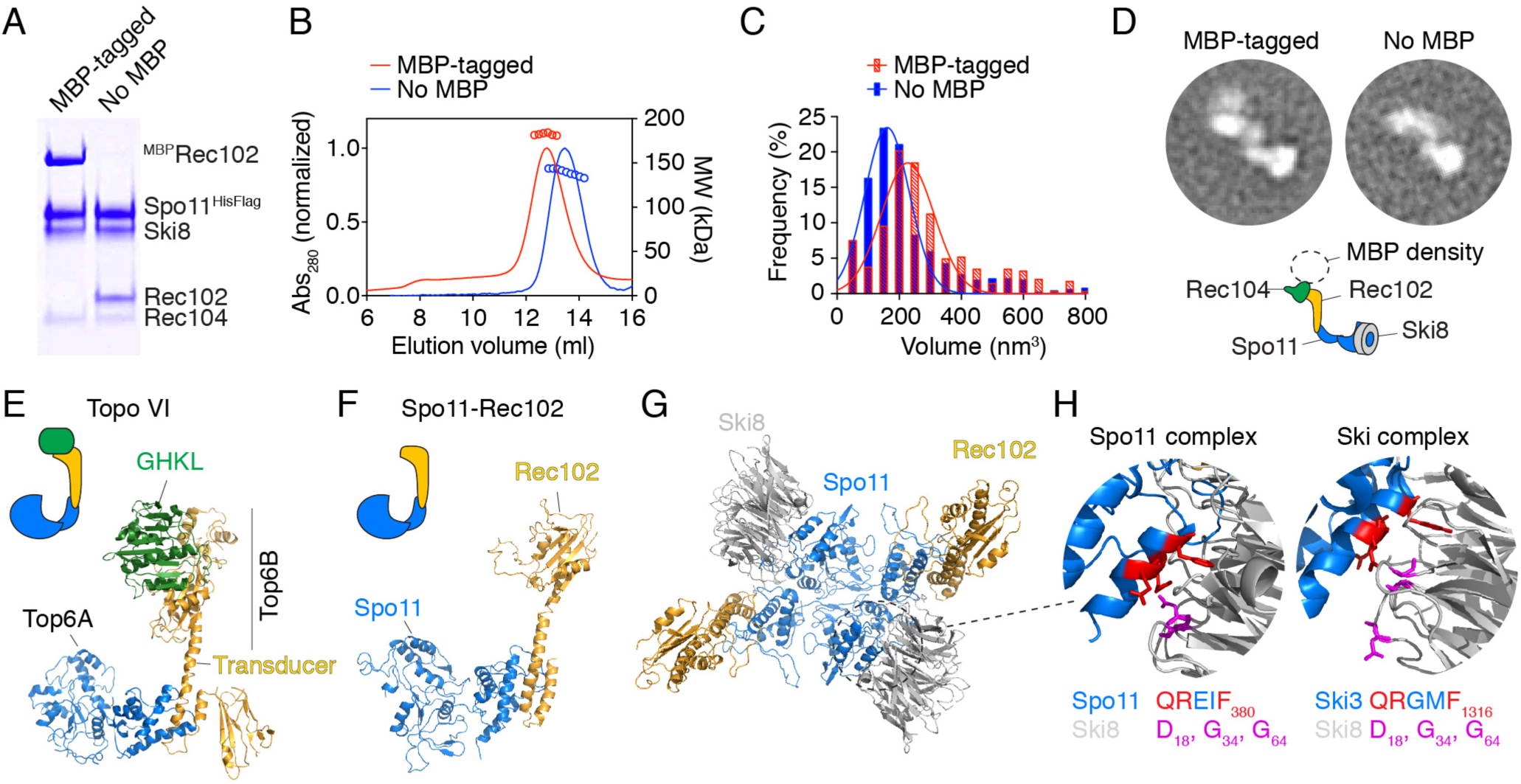
The meiotic DNA double-strand break core complex. A. SDS-PAGE of the purified core complex with and without MBP tag on Rec102 (4 μg per lane). B. SEC-MALS analysis of the core complex with and without MBP tag on Rec102. C. Volumes of core complexes imaged by AFM. D. 2D class averages from nsEM of core complexes with or without a MBP tag on Rec102. Cartoon illustrates subunit positions. E. Structure of Topo VI (Corbett et al., 2007) (PDB: 2Q2E). F. Model of Spo11–Rec102 based on homology with Topo VI (Robert et al., 2016). G. Structural model of a dimer of the Spo11–Rec102–Ski8 complex. Rec104 is not included. H. Interaction between Ski3 and Ski8 within the Ski complex (Halbach et al., 2013) and modeled interaction between Spo11 and Ski8. The motif in Ski3 that interacts with Ski8 (red) is also found in Spo11. Mutation of Q376 in Spo11 abolishes the yeast two-hybrid interaction with Ski8 (Arora et al., 2004) and compromises integrity of the complex (see **Supplementary Figure 3**).

Weight-average molar masses (MW) of complexes with and without MBP (182 kDa and 140 kDa, respectively) determined by size exclusion chromatography followed by multi-angle light scattering (SEC-MALS) were consistent with expected sizes of a heterotetramer (187 kDa and 145 kDa, respectively) (**Figure 1B**). Thus, the purified core complex has a 1:1:1:1 stoichiometry, not 2:2:2:2 as expected based on Topo VI (Corbett et al., 2007; Graille et al., 2008; Robert et al., 2016).

This conclusion was validated by imaging with atomic force microscopy (AFM) and negative-staining electron microscopy (nsEM). The modal volumes of individual particles by AFM were 227 nm^3^ and 161 nm^3^ (with and without MBP, respectively) (**Figure 1C**), consistent with prediction for a heterotetramer (226 nm^3^ and 176 nm^3^, respectively). Two-dimensional class averages of individual particles visualized by EM revealed an elongated shape (**Figure 1D, Supplementary Figure 2**). The MBP-tagged complex displayed additional signal corresponding to only a single MBP subunit, regardless of whether MBP was on Rec102, Rec104, or Spo11 (**Figure 1D, Supplementary Figure S2**).

### A structural model of the core complex

We generated a model of the core complex based on several inputs. Spo11 and Rec102 were previously modeled based on homology with Topo VI (**Figure 1E, F**) (Robert et al., 2016). We constructed a dimeric version representing the putative DSB-competent form (**Figure 1G**), based on the Topo VI holoenzyme (Corbett et al., 2007). Crystal structures of Ski8 have been solved (Cheng et al., 2004; Madrona and Wilson, 2004; Halbach et al., 2013). The interface between Spo11 and Ski8 (**Figure 1H**) was modeled on the Ski complex, within which Ski8 directly contacts a sequence motif in Ski3 that is also present in Spo11 (Halbach et al., 2013). This motif is critical for Spo11–Ski8 interaction (Arora et al., 2004) and for the integrity of the core complex (**Supplementary Figure S3**). The structure and disposition of Rec104 within the complex were completely unknown; we revisit this below.

### Experimental validation of the structural model

We tested aspects of the model using crosslinking coupled to mass spectrometry (XL-MS) (O’Reilly and Rappsilber, 2018) and *in vivo* analysis of site-directed mutants. Samples were treated with the lysine crosslinker disuccinimidyl suberate (DSS) then digested with trypsin and analyzed by mass spectrometry. We identified 861 crosslinks corresponding to 246 distinct pairs of lysines that must be within crosslinkable distance (≤ 27.4 Å for DSS) within the complex (**Figure 2A, Supplementary Table 1**). Mapping Ski8 intramolecular crosslinks onto its crystal structure validated this approach: distances between the α-carbons of all crosslinked lysines were below 27.4 Å (**Supplementary Figure 4A**) and all crosslinked lysines were located away from the predicted interaction surface with Spo11 (**Supplementary Figure 4B**). The XL-MS dataset thus appears appropriately constrained and specific.

**Figure 2:**
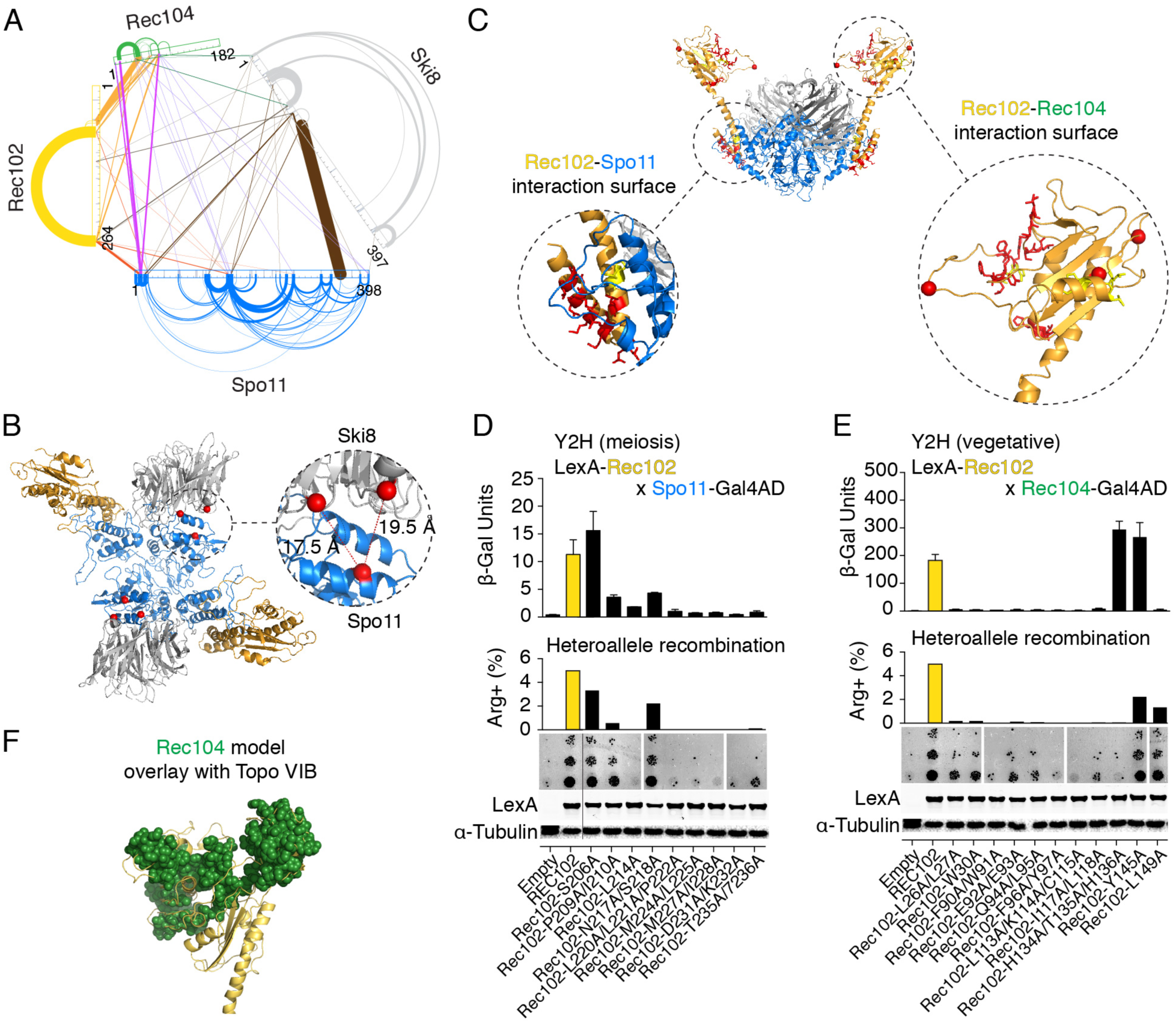
Protein-protein interactions within the core complex. A. XL-MS of the core complex. Arches represent intramolecular crosslinks and lines represent intermolecular crosslinks. Line width is proportional to the number of independent crosslinked peptides and is a proxy for crosslinking frequency. B. Intermolecular crosslinks between Spo11 and Ski8. Distances between α-carbons (red spheres) of crosslinked lysines are shown. C. Positions of mutated residues (red) at predicted interaction surfaces between Rec102 and Spo11 (left), or Rec102 and Rec104 (right). Red spheres are α-carbons of Rec102 lysines that crosslink with Rec104. D, E. Mutagenesis of predicted Rec102 surfaces interacting with Spo11 (D) and Rec104 (E). Top: Yeast-two-hybrid (Y2H) in meiotic or vegetative conditions. Center: *arg4* heteroallele recombination. Bottom: anti-LexA western blotting with α-tubulin as loading control. F. Rec104 model generated by iTasser (spheres) overlaid on Topo VI.

Two intermolecular Spo11–Ski8 crosslinks were particularly abundant, between Spo11 K350 and Ski8 K121 or K147 (fat brown line in **Figure 2A**). These residues lie in close proximity in the model, providing direct experimental support for the inferred arrangement (**Figure 2B**).

The Spo11–Rec102 interface was modeled on the Topo VI holoenzyme structure, in which an N-terminal segment of the A subunit interacts with the C-terminal end of the alpha-helical transducer domain in the B subunit (**Figure 1E, F**) (Corbett et al., 2007; Robert et al., 2016). Potentially consistent with this configuration, we observed intermolecular crosslinks between the Spo11 N-terminus and Rec102 C-terminus (**Figure 2A, Supplementary Table 1**), although the crosslinked residues are not part of the structural model. More importantly, alanine scanning mutagenesis across the predicted interacting region of Rec102 (**Figure 2C**) yielded a series of mutants that had (nearly) completely lost the ability to interact with Spo11 in a yeast two-hybrid assay (see mutations affecting L214 or residues between positions 220 and 236 in **Figure 2D**). All of the interaction-deficient mutants were profoundly defective for meiotic recombination despite being expressed at similar levels as the wild-type protein (**Figure 2D**).

### Rec104 adopts a cryptic GHKL fold

The above findings validate key predictions of our structural model, in turn providing empirical support for the hypothesis that the core complex retains the overall topology expected from its homology with Topo VI. However, important differences are that the B subunit appears truncated such that it lacks a GHKL domain, and a subunit of unknown structure and position (Rec104) is also present.

Based on yeast two-hybrid and other data, Rec102 and Rec104 have been argued to behave as a functional unit (Kee et al., 2004). Rec104 crosslinked efficiently with Rec102 (orange lines in **Figure 2A**), with prominent links mapping to the top of the putative transducer domain of Rec102, at the edge of a five-stranded β-sheet (**Figure 2C**). This position is intriguing because it is where the GHKL domain is seated on the transducer domain in Topo VI (Corbett et al., 2007; Graille et al., 2008) (**Figure 1E**).

To test the importance of this region for the interaction with Rec104, we generated alanine substitutions in Rec102 (**Figure 2C)**. Strikingly, nearly all of these mutations abolished or greatly reduced the interaction with Rec104 and also compromised meiotic recombination (**Figure 2E**; see for example the F90A/W91A double substitution). Exceptions were the triple H134A/T135A/H136A substitution that showed reduced meiotic recombination but normal two-hybrid interaction, the L149A mutant that showed the opposite effect, and the Y145A mutant that behaved like the wild type. None of these mutations appeared to grossly destabilize Rec102, since the mutant proteins were expressed at wild-type levels (**Figure 2E**, bottom). The XL-MS and mutational data thus strongly indicate that Rec104 occupies the position expected for the GHKL domain.

We therefore tested the hypothesis that Rec104 might adopt a cryptic GHKL fold. Remarkably, although secondary structure prediction was uninformative (**Supplementary Figure 5**), iTasser (Yang and Zhang, 2015) generated a three-dimensional model for Rec104 with structural similarity to the GHKL fold even without pre-specifying a template (**Figure 2F**). We thus conclude that Rec104 likely retains the GHKL region of topo VI, albeit a highly diverged, non-catalytic variant.

### DNA bending and binding to duplex junctions

An important biochemical function of Spo11 is to engage its DNA substrate. To gain insights into this interaction, we used AFM to characterize binding of the core complex to linear, relaxed circle, and supercoiled plasmid DNA (**Figure 3A**). This analysis revealed several unanticipated DNA-binding modes.

**Figure 3:**
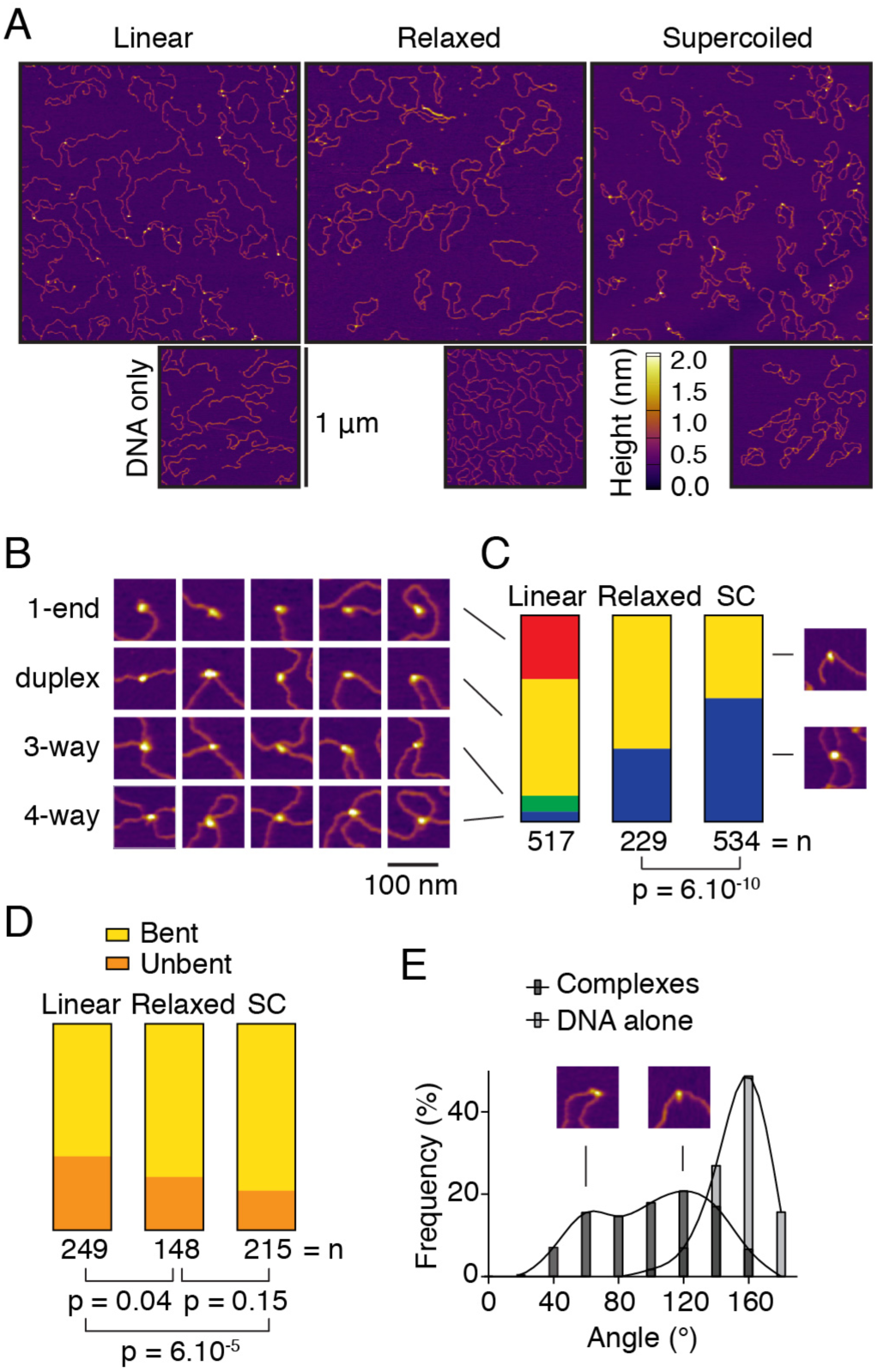
DNA-binding by the core complex analyzed by AFM. A. Wide-field views of core complexes bound to pUC19 plasmids. B. Examples of binding to ends (1-end), internally on duplex DNA (duplex), 3-way junctions, and 4-way junctions. C. Quantification of DNA-bound particles with the three substrates assayed. SC, supercoiled. D. Fractions of duplex-bound particles that exhibited DNA bending. E. Histogram of bending angles (n = 212). Angles at randomly chosen positions along the DNA are shown as a control (n = 115). P values in C, D are from Fisher’s exact test.

With all three DNA forms, 40–65% of the particles were bound internally to the DNA duplex (**Figure 3B, C**), the presumed pre-DSB substrate of Spo11. Particles were frequently associated with DNA bends (**Figure 3D, E**), suggesting that the core complex either bends its substrate or prefers pre-bent DNA. There appeared to be two distinct populations with modal bend angles of ∼60° and ∼120° (**Figure 3E**), possibly representing structural isoforms of the nucleoprotein complexes.

With circular substrates, 35% and 60% of the particles (on relaxed and supercoiled plasmids, respectively) were at a junction between two DNA duplexes (**Figure 3B, C**). Internal duplex binding sites are in substantial molar excess over sites where duplexes happen to cross one another, so we infer that the core complex greatly prefers DNA junctions. This is intriguing because type II topoisomerases, including Topo VI, exhibit DNA-binding specificity for DNA junctions (Corbett et al., 2005; Alonso-Sarduy et al., 2011; Timsit, 2011; Wendorff and Berger, 2018).

### Tight non-covalent interactions with DSB ends

Unexpectedly, one third of protein particles on linear DNA were at an end (**Figure 3B, C**). About 7% of particles were at three-way junctions (**Figure 3B, C**), suggesting simultaneous binding to a DNA end and internally to a duplex. We surmised that Spo11 has intrinsic affinity for the cleavage product, even without a covalent link.

To test this hypothesis, we used electrophoretic mobility shift assays (EMSA) with short oligonucleotide substrates. *In vivo*, DNA cleavage by Spo11 generates a two-base 5′ overhang (Liu et al., 1995; Murakami and Nicolas, 2009; Pan et al., 2011). Using a 25-bp hairpin substrate that carried a “DSB” end that was either blunt or had a 5′-TA overhang, we observed a discrete band of reduced mobility starting at low concentrations of the core complex (≤ 3 nM); higher concentrations yielded an additional slower migrating band (**Figure 4A**). The overhang substrate was bound better than the blunt one, but the shift to the slowest species occurred at similar protein concentrations for both DNA substrates. We interpret the two shifted species as binding to one or both DNA ends with the order of affinities being overhang > blunt > hairpin.

**Figure 4:**
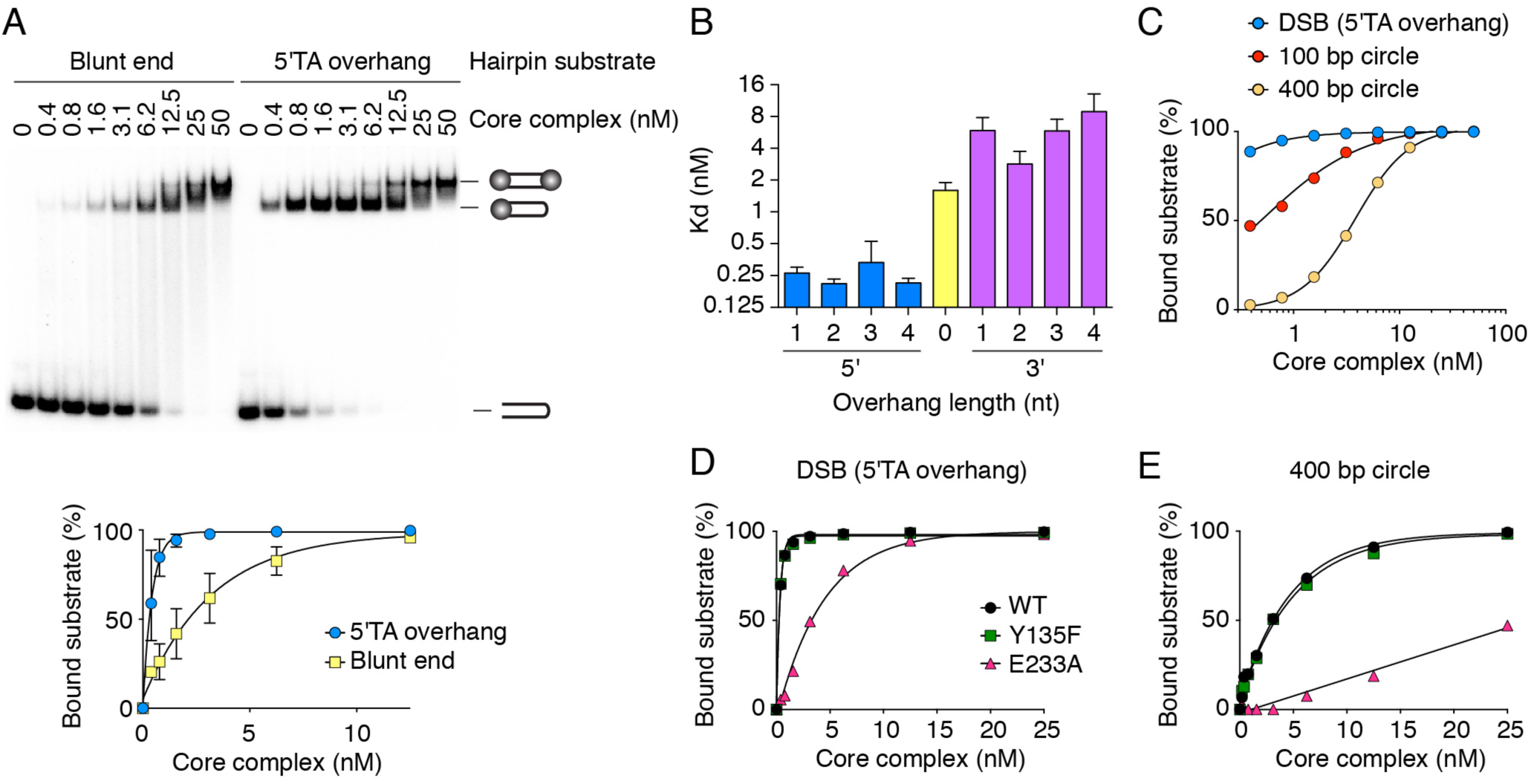
DNA-binding properties of the core complex. A. EMSA of the core complex binding to DNA ends. Core complex was titrated with 5′-labeled 25-bp hairpin substrates with either blunt or two-nucleotide 5′-overhang ends. Quantification (mean ± SD from three experiments) is shown below. B. Apparent dissociation constants for binding (EMSA) to DNA ends with 5′ and 3′ overhangs of different lengths. Error bars are 95% confidence intervals from two experiments. C. Quantification of EMSAs comparing core complex binding to duplex DNA (100-bp or 400-bp DNA circles) vs. DNA ends (25-bp hairpin with 5′-TA overhang). D, E. EMSAs of end binding (D) and duplex binding (E) by Spo11 active-site mutants.

The core complex preferred 5′ overhangs to 3′, with sub-nanomolar apparent dissociation constants (**Figure 4B**). These are remarkably high affinities for non-sequence-specific binding. The core complex preferred a two-base 5′ overhang vs. other lengths in competition experiments (**Supplementary Figure 6A**). End-binding did not require a 5′ phosphate (**Supplementary Figure 6B**).

To compare end binding with binding internally to duplex DNA, we assembled reactions with mini-circles of 100 or 400 base pairs (**Figure 4C**). Compared to the hairpin substrate (K_D_ ≈ 0.1 nM), the mini-circles were bound with lower affinity (K_D_ = 0.5 ± 0.04 nM for 100 bp; 3.8 ± 0.07 nM for 400 bp). We interpret the difference between the circles as reflecting a preference for pre-bent substrates, because the 100 bp circle is more strained and must be highly bent relative to the 400 bp circle (Du et al., 2008). These findings thus agree well with the AFM analysis of plasmid substrates.

Mutating the Spo11 catalytic tyrosine (Y135F) had no effect on binding to the hairpin or the mini-circle, but mutating a presumptive metal-ion-coordinating glutamate (E233A) decreased binding activity ∼10-fold (**Figure 4D, E**). Moreover, presence of Mg^2+^ strongly favored DNA binding (**Supplementary Figure 6C**). These results validate use of the Y135F mutant to compromise catalytic activity while maintaining normal DNA binding *in vivo* (e.g. refs (Cha et al., 2000; Prieler et al., 2005)). Our data may also explain why Y135F can be semi-dominant while E233A is not (Diaz et al., 2002).

### Mapping DNA-binding sites on the core complex

We used FeBABE footprinting to map the DNA-binding surface(s) on the core complex. This method involves conjugation to DNA of a Fe^3+^ ion in an EDTA capsule. Activation with hydrogen peroxide generates hydroxyl radicals that attack nearby proteins. Estimating sizes of cleavage fragments by western blotting of terminally-tagged proteins identifies protein residues that are close to DNA (Miller and Hahn, 2006; Claeys Bouuaert and Keeney, 2017).

Using a 25-bp duplex with a two-base 5′ overhang on one end and a biotin-streptavidin block on the other, we designed two probes that were each conjugated to five FeBABE moieties spread along the twelve base pairs most proximal or distal to the overhang (**Figure 5A**). Binding reactions were analyzed with each of the core complex subunits tagged in turn, all with similar DNA-binding activities (**Supplementary Figure 7**). Hydroxyl radical cleavage yielded specific banding patterns for Spo11, Ski8, and Rec102, and a weak signal for Rec104 (**Figure 5B–E**, lanes 2 and 3). These required presence of the FeBABE probe and were strongly stimulated by hydrogen peroxide (compare lanes 2 and 3 with lanes 1, 4–6). Cleavage of all subunits was more efficient with the overhang-proximal probe than with the distal probe (compare lanes 2 and 3), confirming that the core complex binds to the overhang-containing end.

**Figure 5:**
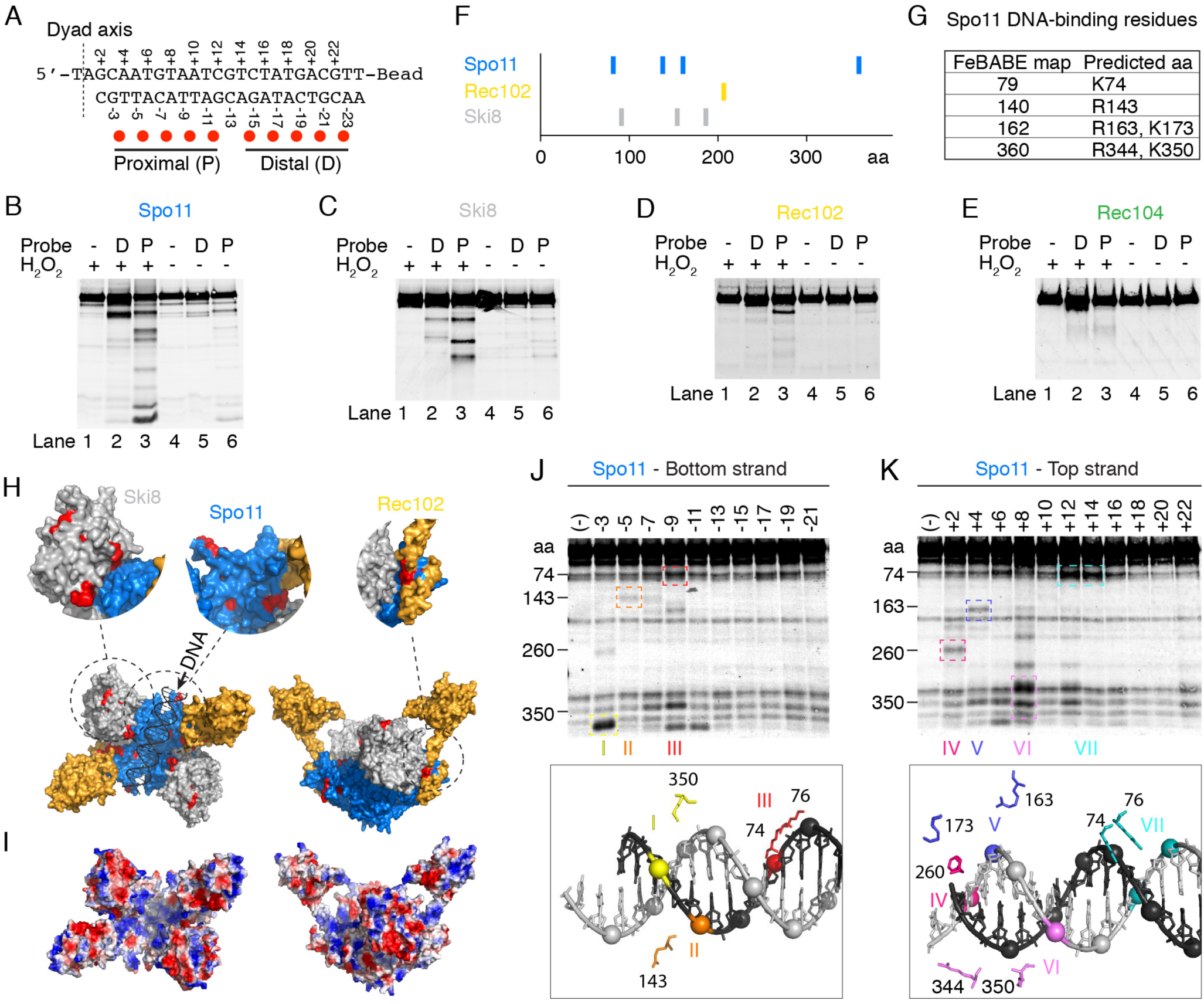
Mapping DNA-binding surfaces by hydroxyl radical footprinting. A. Sequence of the DNA substrate and positions of FeBABE moieties (red dots). The dyad axis is the center of rotational symmetry of Spo11 DSBs. B–E. Hydroxyl radical cleavage of core complexes carrying a C-terminal Flag tag on Spo11 (B), Ski8 (C), or Rec104 (E), or an N-terminal tag on Rec102 (D). Asterisks in panel B indicate cleavage positions illustrated in panel F. F. Summary of hydroxyl radical cleavages. G. Estimated cleavage positions in Spo11 and corresponding predicted DNA-binding residues. H. Hydroxyl radical cleavage sites (red) highlighted on the model of the core complex. I. Electrostatic potential map of the core complex model. J, K. Hydroxyl radical cleavage of tagged Spo11 using probes labeled at single positions along either the bottom (J) or top (K) strands. Prominent cleavage positions are color-coded (roman numerals) and highlighted on the structural model of the end-bound complex (below) to show the spatial correlation with positions of FeBABE-modified phosphates. Some minor cleavage positions were omitted for simplicity. Non-specific degradation fragments were also observed, some of which comigrate with *bone fide* FeBABE-dependent fragments because cleaved positions tend to be surface-exposed.

Estimated cleavage positions and candidate DNA binding (basic) residues are summarized in **Figure 5F, G**, and mapped onto the structural model in **Figure 5H**. Validating the approach, cleavage sites in Spo11 mapped to positions equivalent to the groove that is thought to accommodate DNA in Topo VI (Corbett et al., 2007; Graille et al., 2008). This predicted groove is positively charged in Spo11 (**Figure 5I**).

The main cleavage position in Rec102 (residue ∼207) corresponds to the base of the transducer domain, close to the anticipated DNA-binding groove of Spo11 (**Figure 5H**, top right).

In Ski8, cleavage positions mapped along the surface predicted to point toward the DNA-binding groove of Spo11 (**Figure 5H**, top left). Thus, Ski8 may directly contact DNA in the end-bound complex, in which case the complex must be more closed upon itself than the model based on the Topo VI apoenzyme suggests. Additional support for conformational differences of the end-bound complex is presented below.

To decipher the protein-DNA interactions in finer detail, we examined Spo11-tagged core complexes bound to a set of substrates with a single FeBABE at each of multiple positions on either strand (**Figure 5J, K**). Cleavage fragments were observed when FeBABE was within the first 14 nucleotides from the dyad axis, and each position yielded a distinct cleavage pattern. These findings directly delineate the footprint occupied by the core complex.

We mapped cleavage positions onto a model of Spo11 bound to a DNA end, highlighting those corresponding to specific residues predicted to lie on the surface of the DNA-binding groove (**Figure 5J, K**, bottom panels). The clear spatial correlation between the positions of the FeBABE probe and the predicted positions of DNA-binding residues provides strong experimental support for our model for Spo11 binding to the DSB end.

Based on these results, we altered candidate DNA-interacting residues in Spo11. As predicted, K173A and R344A mutations reduced binding to DNA ends as well as internally to duplexes (**Figure 6A**). *In viv*o, these mutations caused reduced and delayed DSB formation as assessed by labeling of Spo11-oligo complexes (**Figure 6B**) and direct DSB measurements at the *CCT6* hotspot **(Supplementary Figure 8A, B**). Meiotic divisions were also delayed (**Supplementary Figure 8C**), possibly because frequently achiasmate chromosomes trigger the spindle checkpoint (Marston and Wassmann, 2017).

**Figure 6:**
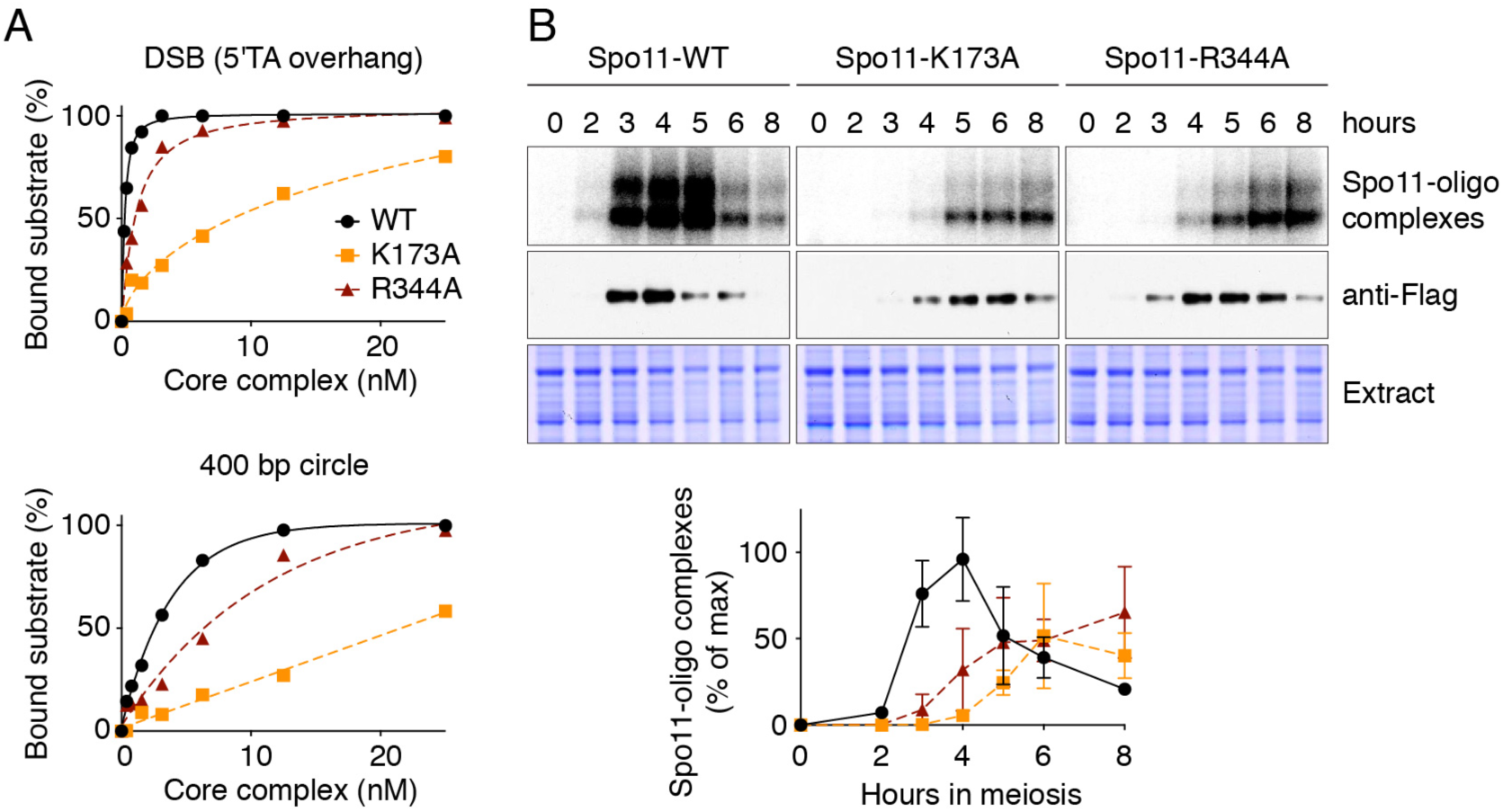
Mutations that affect Spo11 DNA binding compromise DSB formation. A. Binding of wild-type or mutant Spo11-containing core complexes to a 25-bp hairpin substrate with 5′-TA overhang (DSB) or a DNA mini-circle. B. Labeling of Spo11-oligo complexes. Representative gels are shown above; quantification is plotted below (mean and range from two experiments).

### Conformational changes upon DNA binding

XL-MS yielded numerous intramolecular crosslinks within Spo11 (**Figure 2A**). When mapped on our model, at least six pairs of crosslinked residues were further apart than the crosslinker’s 27.4 Å limit (**Figure 7A**). A straightforward interpretation of this incompatibility is that Spo11 might be structurally flexible. Variability in 2D class averages from nsEM supports the interpretation (**Supplementary Figure 2**). The single stretch of polypeptide linking the Toprim and 5Y-CAP domains is a likely flex point (Nichols et al., 1999).

**Figure 7:**
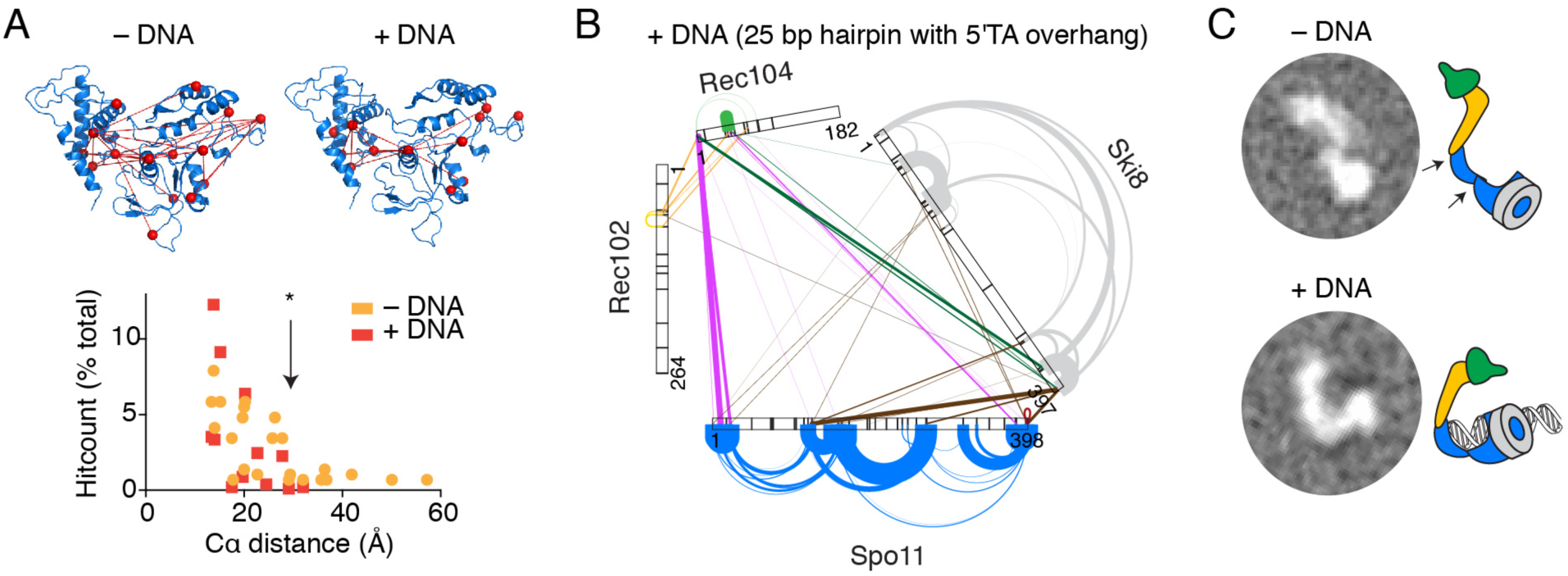
Conformational changes upon DNA binding. A. Model of a Spo11 monomer displaying intramolecular crosslinks without (left) and with (right) DNA. The histogram tabulates distances separating α-carbons of crosslinked lysines. The crosslinkable limit (*) is 27.4 Å. Model-clashing crosslinks would not be explained better by Spo11 dimerization. B. XL-MS of the core complex bound to DNA. C. 2D class averages from nsEM of core complexes in the presence or absence of DNA. DNA substrate in all panels was a 25-bp hairpin with 5′-TA overhang.

These model-clashing intramolecular crosslinks in Spo11 were not observed when XL-MS was performed in the presence of DNA (hairpin substrate with 2-nt 5′ overhang) (**Figure 7A, B**). This supports the interpretation that Spo11 is flexible but adopts a configuration more consistent with the structural model when it binds to DNA.

In addition to this change within Spo11, the presence of DNA substantially changed the crosslinking spectrum across the entire core complex (**Figure 7B**). Noteworthy changes included (i) loss of prominent intramolecular and intermolecular crosslinks involving the C-terminus of Rec102, suggesting that this domain is more constrained in the presence of DNA; (ii) loss of prominent crosslinks between Ski8 and Spo11 (highlighted in **Figure 2B**) presumably because occupancy of the DNA-binding groove of Spo11 obstructs crosslinker access; and (iii) appearance of crosslinks involving the C-terminus of Ski8 with Rec104 and internal parts of Spo11, consistent with the possibility that the end-bound complex adopts a more closed configuration.

These conformational dynamics are also supported by nsEM images, in which the core complex tended to show an elongated shape in the absence of DNA but a more compact, closed configuration in the presence of DNA (**Figure 7C**).

### A DNA-binding mutant of Spo11 alters DSB processing in vivo

To elucidate how the DNA binding activities of Spo11 contribute to its function *in vivo*, we analyzed in detail the phenotype of a binding-defective mutant. We chose F260A, which was originally constructed because the residue was predicted to be in the DNA binding channel (Diaz et al., 2002) (**Figure 8A**). Core complexes containing this mutant bound to DNA with substantially reduced affinity, both at ends (12-fold) and internally (3.2-fold) (**Figure 8B**).

**Figure 8:**
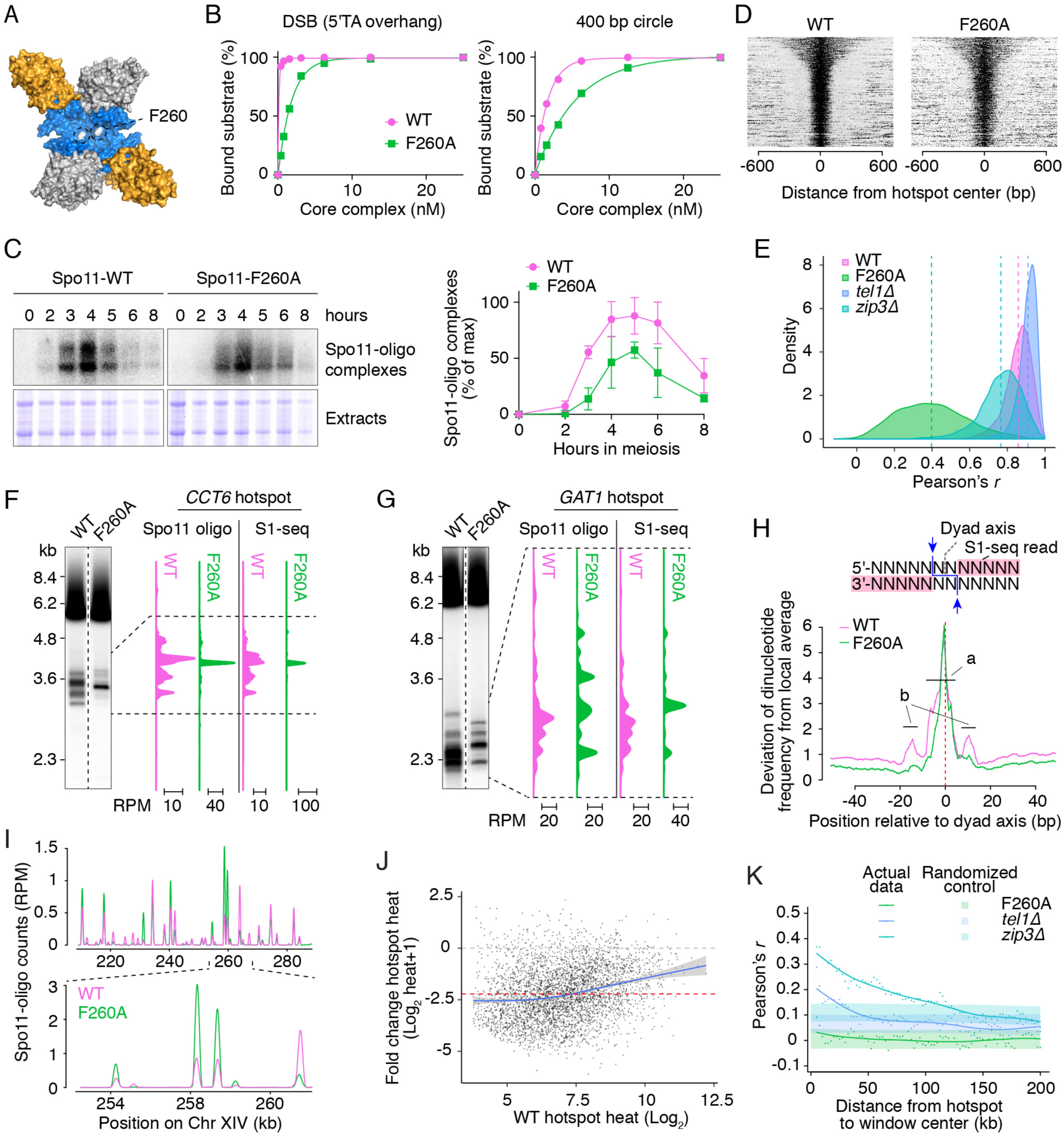
A mutation that affects Spo11-DNA interaction leads to a re-distribution DSBs. A. Position of residue F260 on the core complex. B. Binding of wild-type or F260A mutant core complexes to a 25-bp hairpin substrate with 5′-TA overhang (DSB) or a DNA mini-circle. C. Labeling of Spo11-oligo complexes. Representative gels are shown left, quantifications are plotted right (mean ± SD from three experiments). D. Heatmaps of Spo11 oligos in wild type or F260A at previously defined hotspots (Mohibullah and Keeney, 2017). Hotspots are in decreasing order by width. Spo11-oligo counts for each hotspot were locally normalized to the sum of Spo11 oligos in the window, so color-coding reflects the spatial pattern, not relative signal strength between hotspots. E. Altered distribution of DSBs within hotspots. For each hotspot, we evaluated the correlation (Pearson’s *r*) between the Spo11-oligo pattern in a consensus wild-type map (generated by averaging multiple independent maps (Mohibullah and Keeney, 2017)), and the Spo11-oligo pattern for F260A or the wild-type samples generated in this study. The graph shows the distribution of *r* values from all hotspots (n=3908). Values obtained by comparing consensus wild type to either *tel1Δ* or *zip3Δ* are shown for comparison. F, G. Comparison of Spo11-oligo and S1-seq maps with direct detection of DSBs in *sae2Δ* genomic DNA by Southern blotting at *CCT6* (F) and *GAT1* (G) hotspots. H. Nonrandom dinucleotide composition around Spo11 cleavage sites mapped by S1-seq. At each position, deviation of dinucleotide frequencies from local average was summed. Region a lies within the Spo11 footprint predicted from docking DNA onto the Topo VI structure; regions b lie outside the predicted footprint. The schematic illustrates the positions of Spo11 cleavages (arrows) with respect to the S1-seq read. I. Examples of differences in relative strengths of hotspots between wild type and F260A. Unscaled data in reads per million mapped (RPM) facilitates comparisons of relative strengths between hotspots within a dataset, rather than absolute changes between wild type and mutant. J. Highly variable changes in hotspot strength with Spo11-F260A. Each point is a hotspot defined in wild type (n=3908). Log-fold changes in absolute hotspot strength are plotted as a function of hotspot strength in wild type. The normalized total Spo11-oligo count genome wide in F260A was scaled to 40% of wild type. A Loess trend line is shown in blue. The red line indicates average fold change and the gray dashed line indicates no change. K. Changes in hotspot strength in F260A do not show domains of correlated behavior. Each point compares the log-fold change in hotspots with the change in their neighbors located in a 5-kb window the indicated distance away. Shaded regions show 95% confidence intervals for hotspots randomized within-chromosome. Data for *tel1Δ* (Mohibullah and Keeney, 2017) and *zip3Δ* (Thacker et al., 2014) are shown for comparison.

The F260A mutant had a moderate DSB defect based on analysis at defined genomic locations (Diaz et al., 2002). Quantification of Spo11-oligo complexes extended this genome-wide, indicating that global DSB frequency is reduced to about 40% of wild-type levels based on relative area under the curves for time courses (**Figure 8C**).

Unexpectedly, we also observed a pronounced shift in the size distribution of Spo11 oligos. In wild type, radiolabeled Spo11-oligo complexes migrate on SDS-PAGE as two predominant species that differ by oligo size and that are about equally abundant (Neale et al., 2005), but the F260A mutant gave a strong bias toward shorter oligos (**Figure 8C & Supplementary Figure 9A**). A similar bias was seen with the other DNA binding mutants above (K173A and R344A; **Figure 6B**). These findings suggest that nucleolytic processing of DSBs by Mre11 is influenced by direct DNA binding by Spo11, possibly the noncovalent component of its binding to the products of DNA cleavage. Further implications are addressed in Discussion.

### Spo11 DNA binding shapes the DSB landscape

*S. cerevisiae* DSB distributions are highly nonrandom across chromosomes (Baudat and Nicolas, 1997; Gerton et al., 2000; Blitzblau et al., 2007; Buhler et al., 2007; Pan et al., 2011). Most DSBs form within short regions called hotspots that are typically several hundred base-pairs wide and that mostly coincide with the nucleosome-depleted regions at gene promoters (Wu and Lichten, 1994; Baudat and Nicolas, 1997; Pan et al., 2011; Lam and Keeney, 2015). At broader scales, hotspots occur within large domains (tens of kb) that are relatively DSB-hot or DSB-cold, and at finer scale DSBs are also distributed nonrandomly within hotspots. The factors that shape these distributions over different size scales remain poorly understood (de Massy, 2013; Lam and Keeney, 2015).

We hypothesized that the DNA binding activity of Spo11 influences the finest scale of the DSB landscape by contributing to selection of preferred cleavage sites within hotspots. This idea is supported by previous studies of a few hotspots (Diaz et al., 2002; Murakami and Nicolas, 2009). To test this hypothesis genome-wide, we deep sequenced Spo11 oligos from the F260A mutant and compared to wild type. Because a larger fraction of oligos in the mutant were of the unmapped shorter class, we also mapped unresected DSBs in a *sae2* mutant background using S1-seq. This method prepares DNA ends for sequencing library construction by using S1 nuclease to remove 3′-ssDNA tails of resected DSBs or 5′ overhangs of unresected DSBs (Mimitou et al., 2017; Mimitou and Keeney, 2018). Biological replicate maps were highly reproducible (**Supplementary Figure 9B, C**), so we merged them for further analysis. For results described below, Spo11-oligo mapping and S1-seq were largely indistinguishable.

DSBs in the mutant clustered in hotspots with similar overall dimensions as in wild type (**Figure 8D**), reflecting the constraining influence of the nucleosomes surrounding hotspot nucleosome depleted regions (Pan et al., 2011). However, as predicted, Spo11-F260A generated a dramatically different distribution of DSBs within these hotspots (**Figure 8E**), with similar degrees of difference irrespective of how strong the hotspots were in wild type (**Supplementary Figure 9D**). For example, whereas DSBs were distributed across ∼880 bp of the *CCT6* hotspot in wild type, most of the F260A signal collapsed to a single prominent site which we validated by Southern blotting (**Figure 8F**). Similarly, at the *GAT1* hotspot, most DSBs in the mutant formed at a smaller number of prominent sites than in wild type (**Figure 8G**). Some of the newly prominent sites were also targeted by wild-type Spo11, but often they were either relatively minor sites in wild type or not cleaved at all (e.g., the uppermost band at *GAT1* for the mutant, **Figure 8G**). Thus, Spo11-F260A has substantially altered site selectivity, possibly attributable to its altered affinity for DNA.

Map positions of Spo11 oligos tend to occur in the context of a rotationally symmetric base composition bias that implies preferences of Spo11 for DNA binding and catalysis (Pan et al., 2011). We speculated that the altered site selectivity of F260A would be accompanied by changes in these preferences. To test this, we used the S1-seq maps because these are not subject to a mapping ambiguity caused by rGMP-tailing of Spo11 oligos during library preparation (Pan et al., 2011). In wild-type S1-seq maps, C residues were favored and G residues were disfavored at the position 5′ of the scissile phosphate; and dinucleotides encompassing the scissile phosphate were enriched for 5′-C[A/C/T], TA, AT and AC, but were depleted for GG and GA (**Supplementary Figure 9E, F**). Moreover, there was a ∼20-bp zone of strong sequence bias centered on the dyad axis (“a” region in **Figure 8H**, which corresponds to the predicted footprint deduced by docking Topo VI onto DNA) flanked by sites of weaker bias 11–16 bp on either side of the dyad axis (“b” regions in **Figure 8H**, outside the predicted footprint). These patterns agree well with prior findings from Spo11-oligo sequencing (Pan et al., 2011).

F260A gave dramatically altered patterns, reducing or eliminating most of the normal base composition signatures. There was less enrichment for C and depletion of G 5′ of the scissile phosphate (**Supplementary Figure 9E**) and a corresponding alteration in dinucleotide preferences (i.e., less enrichment (C[A/C/T]) or more depletion (CG) of dinucleotides with 5′-C; **Supplementary Figure 9F**). Other changes within the predicted Spo11 footprint included substantial weakening of the normally strong preference for A at positions –2 to –5 and for T at +2 to +5 (**Supplementary Figure 9E**) and regions outside of the footprint showed a flattening of all of the base composition maps, i.e., substantially less bias (**Figure 8H & Supplementary Figure 9E**). These strong effects of the F260A mutation imply that the intrinsic preference of Spo11 for particular base compositions plays a critical role in cleavage site selection within hotspots.

Unexpectedly, we also observed that relative strengths of hotspots changed substantially in the mutant (**Figure 8I**). To examine these changes, we scaled the F260A map based on the global 2.5-fold decrease in Spo11-oligo complexes (**Figure 8C**) and plotted the absolute log-fold change in the mutant as a function of hotspot strength in wild type (**Figure 8J**). As a point of comparison, we examined similar plots for *tel1Δ* and *zip3Δ* mutants (**Supplementary Figure 9G**). Compared with absence of Tel1 or Zip3, Spo11-F260A caused a wider dispersion of fold changes, and the fold changes showed a modest negative correlation with hotspot strength in wild type. In other words, hotspots vary markedly from one another in their response to F260A, with a tendency that weaker hotspots decrease more. Previous work showed that, in *tel1Δ* and *zip3Δ* mutants, large-scale changes in relative hotspot strength occurred in a domain-wise fashion in which the behavior of individual hotspots was correlated with other hotspots within 50 kb or more (Thacker et al., 2014; Mohibullah and Keeney, 2017). In contrast, hotspot changes in the F260A mutant showed no evidence of local correlation (**Figure 8K**), indicating that the mutation affects DSB frequency in a “hotspot-autonomous” rather than domain-wise fashion. We infer from these results that Spo11 DNA binding activity contributes to a surprising degree to the selection of which hotspot will be cleaved, not just to the selection of cleavage site within a hotspot.

## Discussion

We present a biochemical characterization of the meiotic DSB core complex from *S. cerevisiae* and show that it retains similarities with the molecular architecture and DNA-binding properties of ancestral Topo VI. Nevertheless, key differences stand out, which presumably reflect distinct biological requirements in recombination initiation vs. control of DNA topology.

### Architecture of the core complex

The 1:1:1:1 stoichiometry we observed in solution was unanticipated because Topo VI purifies as a stable A_2_B_2_ dimer of dimers (Corbett et al., 2007; Graille et al., 2008) and because DSB formation requires two Spo11 subunits, one per DNA strand. Mouse SPO11 and TOP6BL were suggested to form 2:2 complexes after expression and purification from *E. coli* (Robert et al., 2016). However, this conclusion was based solely on low-resolution gel filtration and glycerol gradient sedimentation analysis and the recombinant proteins were not shown to retain DNA binding activity, so misfolding and/or nonspecific aggregation cannot be excluded. Thus, while it is possible that the purified yeast and mouse proteins behave differently, the available data are not sufficient to definitively establish the stoichiometry of the mammalian complex. Regardless, we propose that controlled dimerization of yeast Spo11 controls DSB formation *in vivo*. More specifically, we suggest that dimerization of the core complex occurs in the context of its recruitment to the chromosome axis by Rec114. Indeed, previous coimmunoprecipitation experiments indicated that Spo11 self-association *in vivo* depends on the presence of Rec102, Rec104, and Rec114 (Sasanuma et al., 2007).

The core complex retains an overall topology consistent with its relationship with Topo VI. In Topo VI, the GHKL fold within the B subunit has ATPase activity and undergoes ATP-dependent dimerization (Corbett et al., 2007). Dimerization is communicated by the transducer domain to the A subunit to coordinate the formation of a DSB to the passage of an intact duplex through the DNA gate. The mouse and plant TOP6BL subunits retain both GHKL and transducer domains (Robert et al., 2016; Vrielynck et al., 2016). We have provided evidence that, in yeast, the B subunit is split into two polypeptides with Rec102 forming the transducer domain and Rec104 the GHKL domain. Perhaps a similar situation occurs in *S. pombe* and *Drosophila*, where the predicted TOP6BL homolog also lacks the GHKL domain (Bouuaert and Keeney, 2016; Robert et al., 2016).

Although a structural prediction server generated a model for Rec104 with similarity to a GHKL domain, whether Rec104 is truly a degenerate GHKL fold or is rather an unrelated structure that occupies the same position awaits to be clarified. Rec104 does not have any recognizable ATP binding and hydrolysis motifs. The mouse and plant TOP6BL proteins also lack essential ATP-binding residues (Robert et al., 2016; Vrielynck et al., 2016). It has been suggested that topoisomerases use ATP to avoid being trapped in the covalent complex (Bates et al., 2011). Perhaps loss of the ATPase activity of the ancestral topoisomerase provided a straightforward route to yield the meiotic DSB enzyme.

The *S. cerevisiae* core complex also includes Ski8, which has a separate biological function as a part of the Ski complex that provides mRNA to the exosome for degradation (Arora et al., 2004; Halbach et al., 2013). The interaction between Spo11 and Ski8 is essential for DSB activity (Arora et al., 2004) and is required for the integrity of the core complex. However, its function in DSB formation is unclear. Ski8 has a WD40 structure that is often involved in mediating protein-protein interactions (Madrona and Wilson, 2004). It is also possible that Ski8 directly participates in DNA binding. This is supported by the hydroxyl radical footprinting results that showed specific cleavage fragments for Ski8. However, the FeBABE experiment assays for proximity, not interaction, so whether Ski8 directly contacts DNA is unclear. The role of Ski8 in DSB formation is conserved in *S. pombe* and *S. macrospora*, but is not universal because the *Arabidopsis* homolog is not required (Jolivet et al., 2006).

### Substrate specificities and reaction dynamics

AFM analyses revealed different modes of interaction between the core complex and DNA. We propose that these binding modes correspond to intermediates in a multi-step reaction pathway. The data suggest that Spo11 introduces a bend as it engages a DNA duplex, then may trap a second duplex forming a closed dimer encompassing the two duplexes, similar to the mechanism of Topo VI (**Supplementary Figure 10A**) (Corbett et al., 2007; Wendorff and Berger, 2018). In this model, catalysis would occur in the context of a trapped junction, and the core complex then remains tightly associated to the DSB end through non-covalent protein-DNA interactions in addition to the covalent phosphotyrosyl link. The relative frequency at which each binding mode is observed in the AFM experiments suggests that the stability of the complex increases from one intermediate to the next one, which might be expected for a non-catalytic reaction pathway. Nevertheless, our data do not establish the temporal relationship between these complexes. In addition, the stoichiometry of the DNA-bound complexes remains unclear, except for the end-bound complex that is monomeric.

The similarities in DNA-binding properties between the core complex and type II topoisomerases are intriguing. Indeed, the biological function of Spo11 does not require strand passage, so it is unclear why Spo11 retained affinity for DNA junctions (Corbett et al., 2005; Alonso-Sarduy et al., 2011; Timsit, 2011; Wendorff and Berger, 2018). One possibility is that Spo11 captures both sister chromatids as a means to restrict DNA cleavage to just one. Another possibility is that it is related to the tethering the core complex to the chromosome axis: perhaps the uncleaved duplex corresponds to axis DNA while the duplex that experiences the DSB corresponds to the loop. These ideas remain to be tested.

### DNA binding residues and substrate selection by Spo11

Hydroxyl radical footprinting provided the first direct experimental support for the predicted DNA-binding surface within Spo11 and identified residues K173 and R344 as likely DNA-interacting residues that affect cleavage *in vivo*. Genome-wide single-nucleotide resolution DSB mapping with the DNA-binding mutant F260A revealed profound effects on the DSB landscape, showing that Spo11 directly impacts both fine-scale and hotspot-scale selection of cleavage sites.

The attenuated binding affinity of F260A means that Spo11’s preferences contribute less to site selection, and other factors now contribute more, such as chromatin structure (including nucleosome occupancy, H3K4me3 binding by Spp1), competition with other proteins for DNA access, and DNA binding by Spo11-associated proteins. The remaining base composition bias in the mutant is strongly concentrated in the center of the footprint around the dyad axis, suggesting that it reflects direct constraints on catalysis and/or binding.

### The effect of Spo11 DNA binding on DSB processing

The size distribution of Spo11 oligos was affected in the DNA-binding mutants K173A, F260A and R344A, revealing a depletion of the long oligo category. The provenance of the distinct long and short oligo classes was previously unclear (Neale and Keeney, 2009; Pan et al., 2011), but was recently explained through in-depth analyses of Spo11 cleavage patterns (Johnson et al., 2019). It turns out that Spo11 can often cut the same DNA molecule twice in close proximity; when it does, the cuts are spaced at intervals with a 10-bp periodicity that is interpreted to mean that adjacent Spo11 complexes engage the same side of the DNA duplex. This strongly suggests that Spo11 complexes are tethered on a surface (Johnson et al., 2019). Based on the intrinsic property of Rec114–Mei4 and Mer2 to undergo DNA-driven macromolecular condensation (Claeys Bouuaert et al., 2020), we propose that this surface is composed of higher-order Rec114–Mei4– Mer2 nucleoprotein assemblies attached to the chromosome axis (**Supplementary Figure 10B**). This surface is therefore occupied by multiple Spo11 subunits that can simultaneously engage the DNA substrate.

Short oligos are enriched at the edges of DSB hotspots, suggesting that oligo size is dictated by the access of MRX/Sae2, which is proposed to be inhibited at the center of the hotspot by arrays of Spo11 complexes (Johnson et al., 2019). The depletion of the long-oligo class we observed with Spo11 DNA-binding mutants supports this model, because the reduced occupancy of hotspot DNA by the mutant complexes would improve the access of Mre11.

### Spo11 end binding and DSB repair

Usual models envision that Spo11-oligo complexes dissociate from the complementary 3’-end after MRX/Sae2-mediated clipping, although it has been argued that longer oligos might require an active displacement mechanism (Neale et al., 2005; Keeney, 2008). The remarkably tight end-binding activity of the core complex suggests that Spo11-oligo complexes may in fact cap most DSB ends, potentially affecting subsequent repair. In support for this hypothesis, recent mapping of ssDNA–dsDNA junctions in mouse spermatocytes revealed patterns of recombination intermediates consistent with scenarios where the 3’-end remains annealed to the Spo11 oligo (Paiano et al., 2019; Yamada et al., 2019). Since the core complex itself is tethered to the chromosome axis, probably embedded within Rec114–Mei4–Mer2 nucleoprotein condensates, end-capping would therefore maintain a physical connection between the two broken ends during recombination, facilitating their repair and potentially reducing the risks of gross chromosomal rearrangements.

## Materials and Methods

### Preparation of expression plasmids and baculoviruses

Oligos used in this study were purchased from Integrated DNA Technologies. Sequences are listed in **Supplementary Table 2**. Plasmids are listed in **Supplementary Table 3**. Vectors for expression of untagged *S. cerevisiae* Spo11, Ski8, Rec102 and Rec104 were generated by cloning the corresponding open reading frame from SK1 in pFastBac1 (Invitrogen).

*SPO11* was amplified using primers cb753 and cb754, the PCR product and the vector were digested with BamHI and XhoI and cloned to yield pCCB586. *SKI8* was amplified using primers cb755 and cb756, the PRC product and the vector were digested with StuI and XhoI and cloned to generate pCCB587. The open reading frames of *REC102* and *REC104* were cloned into the BamHI and XhoI sites of pFastBac1 to generate pCCB588 and pCCB589.

A sequence coding for a C-terminal 10xHis–3xFlag tag of Spo11 was introduced by inverted PCR and self-ligation using pCCB586 as a template and primers cb757 and cb758 to yield pCCB592. The same strategy was used for C-terminal HisFlag-tagging of Ski8 (template pCCB587, primers cb814 and cb815) to yield pCCB615 and Rec104 (template pCCB589, primers cb816 and cb817) to yield pCCB616. N-terminal HisFlag-tagging of Rec102 was achieved by cloning Rec102 in pFastBac-HTb-Flag, which codes for 6xHis and 2xFlag tags, to yield pCCB633.

N-terminal MBP-tagged vectors were generated by amplifying the MBP gene from pMAL-c2x and cloning into the untagged expression construct using appropriate restriction enzymes. The primers used for amplifying the MBP gene were the following (with resultant vector between brackets): Spo11: cb830 and cb831 (pCCB620), Ski8: cb820 and cb821 (pCCB618), Rec102: cb818 and cb819 (pCCB617), Rec104: cb822 and cb823 (pCCB619).

Point mutants were generated by QuikChange mutagenesis. HisFlag-tagged Spo11 mutants used pCCB592 as a template. Plasmid number and mutagenesis primers for each mutant are as follows: Spo11-Y135F (cb806 and cb807, pCCB611), Spo11-E233A (cb810 and cb811, pCCB613). Spo11-F260A (cb896 and cb897, pCCB646), Spo11-K173A, (cb1299 and cb1230, 814), Spo11-R344A (cb1301 and cb1302, pCCB815). The viruses were produced by a Bac-to-Bac Baculovirus Expression System (Invitrogen) following the manufacturer’s instructions.

The yeast two-hybrid vector expressing the LexA-Rec102 fusion (pSK282) was reported previously (Maleki et al., 2007). Rec102 mutants were generated by Quikchange mutagenesis using pSK282 as a template. Mutagenesis primers and resultant plasmids are as follows: Rec102-L26A/L27A (cb834 and cb835, pCCB655), Rec102-W30A (cb836 and cb837, pCCB656), Rec102-F90A/W91A (cb838 and cb839, pCCB657), Rec102-E92A/E93A (cb840 and cb841, pCCB658), Rec102-Q94A/L95A (cb842 and cb843, pCCB659), Rec102-F96A/Y97A (cb844 and cb845, pCCB660), Rec102-L113A/K114A/C115A (cb846 and cb847, pCCB661), Rec102-I117A/L118A (cb848 and cb849, pCCB662), Rec102-H134A/T135A/H136A (cb850 and cb851, pCCB663), Rec102-Y145A (cb852 and cb853, pCCB664), Rec102-L149A (cb854 and cb855, pCCB665), Rec102-S206A (cb856 and cb857, pCCB666), Rec102-P209A/I210A (cb858 and cb859, pCCB667), Rec102-L214A (cb860 and cb861, pCCB668), Rec102-N217A/S218A (cb862 and cb863, pCCB669), Rec102-L220A/L221A/P222A (cb864 and cb865, pCCB670), Rec102-M224A/L225A (cb866 and cb867, pCCB671), Rec102-M227A/I228A (cb868 and cb869, pCCB672), Rec102-D231A/K232A (cb870 and cb871, pCCB673), Rec102-T235A/T236A (cb872 and cb873, pCCB674). All the mutant Rec102 vectors were verified by sequencing using primer cb902. The Gal4AD-Spo11 (pSK305) and Gal4AD-Rec104 (pSK310), empty Gal4AD (pSK276) and LexA (pSK272) vectors have been described previously (Arora et al., 2004).

### Expression and purification of recombinant proteins

Expression of the Spo11 core complexes was achieved by co-infection of *Spodoptera frugiperda* Sf9 cells with an appropriate combination of viruses at a multiplicity of infection (MOI) of 2.5 each. Spo11^HisFlag^:Ski8:Rec102:Rec104 complex used viruses produced from the vectors pCCB592, pCCB587, pCCB588 and pCCB589. Spo11^HisFlag^:Ski8:^MBP^Rec102:Rec104 complex used viruses produced from the vectors pCCB592, pCCB587, pCCB617 and pCCB589. Other tagged or mutant complexes were prepared using the appropriate combination of viruses (see list of vectors in **Supplementary Table 3**).

Cells were harvested 62 h after infection, washed with phosphate buffer saline (PBS), frozen in dry ice and kept at -80 °C until use. A typical purification was performed with cell pellets from 500 ml to 2 l culture. All the purification steps were carried out at 0-4 °C. Cell pellets were resuspended in 4 volumes of lysis buffer (25 mM HEPES-NaOH pH 7.5, 500 mM NaCl, 0.1 mM DTT, 20 mM imidazole, 1× Complete protease inhibitor tablet (Roche) and 0.1 mM phenylmethanesulfonyl fluoride (PMSF)). Cells were lysed by sonication and centrifuged at 43,000 g for 30 min. The cleared extract was loaded onto 1 ml pre-equilibrated NiNTA resin (Qiagen). The column was washed extensively with Nickel buffer (25 mM HEPES-NaOH pH 7.5, 500 mM NaCl, 10% glycerol, 0.1 mM DTT, 20 mM imidazole, 0.1 mM PMSF). The tagged complexes were then eluted in Nickel buffer containing 250 mM imidazole. MBP-tagged complexes were further purified on amylose resin (NEB). The core complex without MBP was instead further purified on Anti-Flag M2 affinity resin (Sigma). Fractions containing protein were pooled and diluted in 3 volumes of Amylose/Flag buffer (25 mM HEPES-NaOH pH 7.5, 500 mM NaCl, 10% glycerol, 2 mM DTT, 5 mM EDTA). Next, the complexes were bound to 1 ml of the appropriate resin in a poly-prep chromatography column (Bio-Rad) and the resin was washed extensively with buffer. Complexes were eluted from amylose resin with buffer containing 10 mM maltose, or from anti-Flag resin with buffer containing 250 μg/ml 3xFlag peptide (Sigma). Fractions containing protein were pooled and loaded on a Superdex 200 column preequilibrated in Amylose/Flag buffer. Fractions containing protein were concentrated in 50-kDa-cutoff Amicon centrifugal filters (Millipore). Aliquots were frozen in dry ice and stored at -80 °C. Mutant core complexes were prepared using the same procedure as the wild-type proteins.

### Substrates for DNA-binding assays

DNA substrates were generated by annealing complementary oligos (sequences in **Supplementary Table 2**). Oligos over 40 nucleotides were first purified on 10% polyacrylamide-urea gels. They were subsequently mixed in equimolar concentrations (typically 10 μM) in STE (100 mM NaCl, 10 mM Tris-HCl pH 8, 1 mM EDTA), heated and slowly cooled on a PCR thermocycler (98 °C for 3 min, 75 °C for 1 h, 65 °C for 1 h, 37 °C for 30 min, 25 °C for 10 min).

Hairpin substrates for EMSA were assembled by self-annealing of the following primer: blunt end (cb953); 5′-overhang of 1 nt (cb1061), 2 nt (cb957), 3 nt (cb1062), 4 nt (cb1063); 3′-overhang of 1 nt (cb1064), 2 nt (cb1065), 3 nt (cb1066), 4 nt (cb1067). Substrates were 5′ end-labeled with [γ-^32^P]-ATP (Perkin Elmer) and T4 polynucleotide kinase (NEB) and labeled substrates were purified by native polyacrylamide gel electrophoresis. The 100-bp mini-circle was prepared as follows: oligos cb1027 and cb1028 were labeled with [γ-^32^P]-ATP (Perkin Elmer) by T4 polynucleotide kinase. Labeled oligos were mixed and annealed by boiling and slow cooling. The mini-circle was then ligated with T4 DNA ligase and the circle substrate was purified by native polyacrylamide gel electrophoresis. The 400-bp mini-circle was prepared by PCR amplification of pUC19 plasmid DNA with phosphorylated primers cb962 and cb963 in the presence of α-^32^P-dCTP and self-ligation in dilute conditions by T4 DNA ligase. The DNA was concentrated by ethanol precipitation and the mini-circle purified by native agarose electrophoresis.

Substrates for hydroxyl radical footprinting used the 3′-biotinylated oligo cb958 and either the unmodified oligo cb959 (5’-TA-control) or the phosphorotioate-containing oligo cb560 (5′-TA-distal) or cb561 (5′-TA-proximal). FeBABE probes in **Figure 5J** used oligo cb598 annealed with cb1072 to cb1081. FeBABE probes in **Figure 5K** used oligo cb1082 annealed with cb1083 to cb1093. Oligos were annealed in 40 μl STE at a concentration of 5 μM biotinylated oligos and a 1.2-fold excess of non-biotinylated oligos, then ethanol precipitated and resuspended in 20 mM MOPS pH 7.9. FeBABE conjugation to phosphorothioate-containing DNA substrates were in 20 μl reactions in 20 mM MOPS (pH 7.9) containing 4 μM DNA and 3.5 mM FeBABE (Dojindo). After 16 h incubation at 50 °C, the substrates were immobilized to 800 ng M280 streptavidin-coated Dynabeads (Invitrogen) in 400 μl 20 mM MOPS (pH 7.9) for 4 hours at 4 °C. Excess FeBABE was removed by washing three times with 500 μl 20 mM MOPS pH 7.9 and one time with 200 μl storage buffer (25 mM Tris-HCl pH 7.5, 10% glycerol, 100 mM NaCl, 200 μg/ml BSA). Substrates were resuspended in storage buffer at an estimated concentration of 1000 pmol/μl beads (final substrate concentration of about 1 μM) and stored at 4 °C.

### Electrophoretic mobility shift assays

Binding reactions (20 μl) were carried out in 25 mM Tris-HCl pH 7.5, 7.5% glycerol, 100 mM NaCl, 2 mM DTT, 5 mM MgCl_2_ and 1 mg/ml BSA with 0.1 nM DNA (for hairpin substrates) or 1 nM DNA (for mini-circles) and the indicated amounts of protein complexes. Complexes were assembled for 30 minutes at 30 °C and separated on a 5% Tris-acetate-polyacrylamide/bis (80:1) gel containing 0.5 mM MgCl_2_ at 200 V for two hours. Gels were dried, exposed to autoradiography plates and revealed by phosphorimaging (Fuji).

### Hydroxyl radical cleavage assay

Binding reactions (20 μl) were carried out in 25 mM Tris-HCl pH 7.5, 10% glycerol, 100 mM NaCl, 1 mg/ml BSA, 5 mM MgCl_2_ with 25 nM immobilized substrates and 25 nM Spo11 core complex. Complexes were assembled for 10 minutes at 30 °C, washed twice with 200 μl FeBABE buffer (25 mM Tris-HCl pH 7.5, 10% glycerol, 100 mM NaCl, 5 mM MgCl_2_, 0.01% NP-40) with 2 mM DTT and once with 200 μl buffer without DTT. Reactions were resuspended in 20 μl FeBABE buffer. One half (10 μl) was treated with 1.25 μl of 50 mM sodium ascorbate followed rapidly with 1.25 μl of 50 mM H_2_O_2_ in 10 mM EDTA and the cleavage reaction performed at 30 °C for 10 minutes. The other half was left untreated as a negative control. Reactions were quenched with 6 μl 4× LDS sample buffer and 1 μl 1 M DTT, boiled for 5 minutes and loaded on 4– 12% NuPAGE Bis-Tris gels (to analyze the Spo11 and Ski8 fragments) or 10% gels (to analyze the Rec102 and Rec104 fragments) in MES running buffer. After electrophoresis, proteins were transferred onto Immobilon-FL PVDF membranes (Millipore) and membranes were blotted with Anti-Flag M2 antibody (Sigma) followed by a IRDye 680RD Goat anti-mouse IgG (Li-COR). Western blots were revealed using the Li-COR Bioscience Odyssey infrared imaging system.

### Mapping of the protein-DNA interface

The molecular sizes of Flag-tagged fragments of Spo11, Ski8, Rec102 and Rec104 generated by the hydroxyl radical cleavage assay were determined by comparing their migration with MagicMark™ XP Western protein standards (Invitrogen). Profiles of each lane were quantified using ImageGauge. The peaks were identified for each molecular weight standard and used to deduce a fourth order polynomial equation of the migration distance as a function of molecular weight. The equation was used to deduce the molecular weights of the protein fragments, which in turn provide an estimate position of the hydroxyl radical cleavage sites. The accuracy with which cleavage fragments are estimated was assessed by comparing the calculated cleavage sites of trypsin and chymotrypsin digestions with theoretical cleavage sites. Based on multiple experiments, we estimate that the cleavage sites are mapped within 5–10 residues.

### Structural modeling

The structure of the Ski complex is from PDB accession number 4BUJ (Halbach et al., 2013). Intramolecular crosslinks on Ski8 were mapped on 1SQ9 (Madrona and Wilson, 2004). The homology-based model for Spo11 and Rec102 was reported previously (Robert et al., 2016). Secondary structure predictions were generated by PsiPred (McGuffin et al., 2000). The 3D model of Rec104 was generated by iTasser (Yang and Zhang, 2015). The model was generated without a template to guide the modeling and picked up Topo VI structures (PDB: 1MU5 and 2Q2E) as the closest matches. The model was generated with the following scores: C-score = -2.15, estimated TM-score = 0.46 ± 0.15, estimated RMSD = 10.0 ± 4.6 Å. The model of the core complex was generated using Pymol. The pdb files are available upon request.

### AFM imaging

Relaxed and linear plasmid substrates for AFM imaging were prepared by treatment of pUC19 with topoisomerase I and NdeI, respectively. Core complexes were diluted to a final concentration of 4 nM in the presence of 1 nM DNA (supercoiled, relaxed or linear) in 25 mM HEPES-NaOH pH 6.8, 5 mM MgCl_2_, 50 mM NaCl, 10% glycerol. Complexes were assembled at 30 °C for 30 minutes. A volume of 40 µl of the protein-DNA binding reaction was deposited onto freshly cleaved mica (SP1) for 2 minutes. The sample was rinsed with 10 ml ultrapure deionized water and the surface was dried using a stream of nitrogen. AFM images were captured using an Asylum Research MFP-3D-BIO (Oxford Instruments) microscope in tapping mode at room temperature. An Olympus AC240TS-R3 AFM probe with resonance frequencies of approximately 70 kHz and spring constant of approximately 1.7 N/m was used for imaging. Images were collected at a speed of 0.5–1 Hz with an image size of 2 µm at 2048 × 2048 pixel resolution.

For AFM imaging of proteins alone, samples were diluted in 25 mM HEPES-NaOH pH 7.5, 500 mM NaCl, 10% glycerol, 2 mM DTT, 5 mM EDTA. Immobilized proteins were imaged with a Bruker FMV-A AFM probe with resonance frequencies of approximately 75 kHz and spring constant of approximately 2.5 N/m. Images were collected at a speed of 0.8 Hz with an image size of 1 µm at 512 × 512 pixel resolution. For volume analyses, raw data were exported into 8-bit grayscale tiff images using the Asylum Research’s Igor Pro software. Tiff images were then imported into FIJI/ImageJ (NIH) for quantification of volume using a custom written FIJI code. A height threshold of 0.25 nm was set for each protein sample, as well as a pixel area threshold, to exclude noise from the image. The volume of each structure was calculated using the formula *V = I*_*avg*_ · *Z*_*conversion*_ · *A*_*p*_ · *XY*_*conversion*_^*2*^ where *I*_*avg*_ is the average intensity, *Z*_*conversion*_ is the conversion of one gray scale unit of intensity into height in nanometers, *XY*_*conversion*_ is the pixel to nanometer conversion for the image in xy, and *A*_*p*_ is the area of particles in pixels. The predicted volume of protein complexes can be calculated according to the equation: V_c_ = V_1_ × M_0_ / N_0_, where M_0_ is the molecular weight, N_0_ is Avogadro’s number and V_1_ is the specific volume for proteins (0.73 cm^3^ g^-1^) (Richards, 1985). Based on this equation, the predicted volume of a 1:1:1:1 core complex with MBP tag (187.8 kDa) or without MBP tag (145.3 kDa) is 226 nm^3^ and 176 nm^3^, respectively.

Additional FIJI analysis was done for the quantification of DNA bending angles at protein-DNA interaction sites. A custom-written FIJI macro was written to allow for two lines to be manually drawn on DNA strands on either side of a DNA-protein binding event to allow for automated calculation of the angles created by the DNA strand.

### Yeast strains and targeting vectors

Yeast strains were from the SK1 background. All strains used in this study are listed in **Supplementary Table 4**. Flag-tagged Spo11 strains were constructed by transformation of Sph1 fragments of plasmids pSK806 (WT), pSK809 (F260A), pCCB822 (K173A) and pCCB823 (R344A), replacing the endogenous locus with the tagged construct together with a hygromycin resistance cassette.

### Crosslinking – mass spectrometry

For protein-alone samples, ∼25 µg protein complexes were cross linked in 160 µl reactions in the presence of 2 mM disuccinimidyl suberate (DSS) in buffer containing 25 mM HEPES-NaOH pH 7.5, 500 mM NaCl, 10% glycerol, 2 mM DTT, 5 mM EDTA. After 10–20 minutes at 30 °C, crosslinking reactions were quenched with 100 mM Tris-HCl pH 7.5. For crosslinking reactions in the presence of DNA, 20 µg core complexes was bound to equimolar concentrations (100 nM) of a 25 bp hairpin substrate with 2 nt 5′-overhang in a 6-ml reaction volume. Binding reactions and crosslinking were performed as above, but with 100 mM NaCl and 5 mM MgCl_2_ (instead of EDTA). After crosslinking, proteins were concentrated by acetone precipitation and resuspended in 1× Laemmli buffer. Samples were boiled, separated by SDS-polyacrylamide gel electrophoresis and stained with SimplyBlue SafeStain (Invitrogen). Slower-migrating bands representing protein-protein crosslinks were excised and digested *in situ* with trypsin digestion as described (Sebastiaan Winkler et al., 2002). The tryptic peptides were purified using a 2-µl bed volume of Poros 50 R2 (Applied Biosystems) reversed-phase beads packed in Eppendorf gel-loading tips (Erdjument-Bromage et al., 1998). The digested peptides were diluted to 0.1% formic acid, and each sample was analyzed separately by microcapillary LC with tandem MS by using the NanoAcquity system (Waters) with a 100 µm inner diameter × 10 cm length C18 column (1.7 µm BEH130; Waters) configured with a 180 µm × 2 cm trap column coupled to a Q-Exactive Plus mass spectrometer (Thermo Fisher Scientific). A proxeon nanoelectrospray source set at 1800 V and a 75 µm (with 10 µm orifice) fused silica nano-electrospray needle (New Objective, Woburn, MA) was used to complete the interface. 1 µl of sample was loaded onto the trap column, washed with 3× loop volume of buffer A (0.1% formic acid) and the flow was reversed through the trap column and the peptides eluted with a 1–50% acetonitrile (with 0.1% formic acid) gradient over 50 min at a flow rate of 300 nl/min over the analytical column. The QE Plus was operated in automatic, data-dependent MS/MS acquisition mode with one MS full scan (370–1700 *m*/*z*) at 70,000 mass resolution and up to ten concurrent MS/MS scans for the ten most intense peaks selected from each survey scan. Survey scans were acquired in profile mode and MS/MS scans were acquired in centroid mode at 17500 resolution and isolation window of 1.5 amu. AGC was set to 1 × 10^6^ for MS1 and 5 × 10^5^ and 100 ms maximum IT for MS2. Charge exclusion of 1, 2 and greater than 8 enabled with dynamic exclusion of 15 s. To analyze the cross-linked peptides we used pLink (Yang et al., 2012). The raw MS data was analyzed using pLink search with the following parameters: precursor mass tolerance 50 p.p.m., fragment mass tolerance 10 p.p.m., cross-linker DSS (cross-linking sites K and protein N terminus, xlink mass-shift 138.068, monolink mass-shift 156.079), fixed modification C 57.02146, variable modification oxidized methionine, deamidation N,Q, protein N-acetyl, peptide length minimum 4 amino acids and maximum 100 amino acids per chain, peptide mass minimum 400 and maximum 10,000 Da per chain, enzyme trypsin, two missed cleavage sites per chain (four per cross-link). The data were imported on the xiNET online tool to generate crosslinking maps (Combe et al., 2015). All the identified crosslinks can be found in **Supplementary Table 1**.

### SEC-MALS

The light scattering data were collected using a Superdex 200, 10/300, HR Size Exclusion Chromatography (SEC) column (GE Healthcare, Piscataway, NJ), connected to High Performance Liquid Chromatography System (HPLC), Agilent 1200, (Agilent Technologies, Wilmington, DE) equipped with an autosampler. The elution from SEC was monitored by a photodiode array (PDA) UV/VIS detector (Agilent Technologies, Wilmington, DE), differential refractometer (OPTI-Lab rEx Wyatt Corp., Santa Barbara, CA), static and dynamic, multiangle laser light scattering (LS) detector (HELEOS II with QELS capability, Wyatt Corp., Santa Barbara, CA). The SEC-UV/LS/RI system was equilibrated in buffer 25 mM Hepes pH 7.5, 500 mM NaCl, 10% glycerol, 2 mM EDTA at the flow rate of 0.5 ml/min or 1.0 ml/min. Two software packages were used for data collection and analysis: the Chemstation software (Agilent Technologies, Wilmington, DE) controlled the HPLC operation and data collection from the multi-wavelength UV/VIS detector, while the ASTRA software (Wyatt Corp., Santa Barbara, CA) collected data from the refractive index detector, the light scattering detectors, and recorded the UV trace at 280 nm sent from the PDA detector. The weight average molecular masses were determined across the entire elution profile in the intervals of 1 sec from static LS measurement using ASTRA software as previously described (Folta-Stogniew and Williams, 1999).

### Negative staining EM

The sample (4 µl of protein in 25 mM HEPES-NaOH pH 7.5, 500 mM NaCl, 10% glycerol, 2 mM DTT, 5 mM EDTA) was applied to fresh, glow-discharged homemade continuous carbon grids. For the DNA-bound sample, the complex was mixed with a 25-bp hairpin substrate with 5′-TA overhang in 25 mM HEPES-NaOH pH 7.5, 100 mM NaCl, 10% glycerol, 2 mM DTT, 2 mM MgCl_2_. After 1-minute incubation, the grids were stained in five consecutive 40-µl drops of 2% uranyl acetate during a total of 45 seconds prior to blotting completely dry. Negatively-stained specimens were examined under an FEI Tecnai-12 TWIN electron microscope equipped with a LaB_6_ filament and operated at 100 kV acceleration voltage. Data were acquired using a dose of ∼25 e^-^/Å^2^ at a nominal magnification of 49,000 × and binned to 4.14 Å per pixel. All images were recorded on a FEI Eagle 4k × 4k pixel camera (Thermo) utilizing LEGINON data collection software (Suloway et al., 2005). Particle picking, contrast transfer function estimation and phase-flipping correction were performed using EMAN (Ludtke et al., 1999; Tang et al., 2007). Reference-free 2D methods implemented in EMAN (Ludtke et al., 1999), and 2D maximum-likelihood classification as implemented in RELION (Scheres, 2012), were used to analyze an average of 10,000 particles for each condition. The particles were too conformationally heterogeneous to permit 3D volume reconstruction.

### Spo11-oligo labeling

The procedure for labeling Spo11-oligo complexes has been described (Neale and Keeney, 2009). Briefly, yeast cultures were harvested at the indicated time into meiosis and denatured extracts were prepared by trichloroacetic acid precipitation. Proteins were solubilized in 2% SDS, 500 mM Tris-HCl pH 8.1, 10 mM EDTA. Extracts were diluted in an equal volume of 2× IP Buffer (2% Triton X100, 30 mM Tris-HCl pH 8.1, 300 mM NaCl, 2 mM EDTA, 0.02% SDS) and Flag-tagged Spo11-oligo complexes were immunoprecipitated on agarose beads conjugated with mouse monoclonal M2 anti-Flag antibody. DNA was labeled on the beads with terminal deoxynucleotidyl transferase and [α-^32^P]-dCTP. After washing the beads in 1× IP buffer, proteins were eluted with LDS sample buffer and separated by SDS-PAGE. The gel was dried and developed by autoradiography. For analysis of Spo11-oligo length, radiolabeled Spo11-oligo complexes were digested with proteinase K and deproteinized oligos were separated on a 15% TBE-UREA polyacrylamide gel.

### Southern blot analysis of meiotic DSB formation

Meiotic DSB analysis by Southern blotting was performed as described (Murakami et al., 2009). Briefly, synchronized cultures undergoing meiosis were harvested at the indicated times. After DNA purification, 800 ng of genomic DNA was digested with PstI and separated on a 1% TBE-agarose gel. For direct comparison of break formation at *GAT1* and *CCT6* with genome-wide DSB maps, genomic DNA from *sae2* strains were prepared in agarose plugs as described (Murakami et al., 2009). Plugs were equilibrated in NEB CutSmart buffer, digested with PstI-HF, and digested DNA was separated on a 0.65% TBE-agarose gel. DNA was transferred to Hybond-XL nylon membranes by vacuum transfer, hybridized with *GAT1* probe (amplified with primers 5′-CGCGCTTCACATAATGCTTCTGG and 5′-TTCAGATTCAACCAATCCAGGCTC) or *CCT6* probe (amplified with primers 5′-GCGTCCCGCAAGGACATTAG and 5′-TTGTGGCTAATGGTTTTGCGGTG), and developed by autoradiography.

### Heteroallele recombination assays

Heteroallele recombination assays were done in diploid strains carrying the *arg4-Bgl* and *arg4-Nsp* alleles (Nicolas et al., 1989). Three independent colonies were cultured overnight in selective dropout medium, then transferred to YPA for 13.5 hours prior to meiotic induction in 2% KOAc. After at least 6 hours in meiosis, appropriate dilutions of cultures were plated on arginine dropout medium to measure frequency of Arg+ recombinants and on YPD to measure colony forming units.

### Yeast extracts and western blotting

Denaturing whole-cell extracts were prepared in10% trichloroacetic acid by agitation in the presence of glass beads. Precipitated proteins were solubilized in Laemmli sample buffer and appropriate amounts of protein were separated by SDS-PAGE and analyzed by western blotting. Antibodies for western blotting were goat polyclonal anti-LexA (1:2000, Santa Cruz), rat polyclonal anti-α-tubulin (1:5000, Bio-rad), mouse monoclonal anti-Flag M2 (1:2000, Sigma) HRP-conjugated mouse monoclonal anti-Flag M2 (1:2000, Sigma), mouse monoclonal anti-MBP (1:2000, NEB), rabbit polyclonal anti-Spo11 (1:1000, this laboratory). Secondary antibodies were used at 1:5000 dilution: IRDye 800CW goat anti-mouse IgG (LI-COR), HRP-conjugated goat anti-mouse IgG (Bio-Rad), HRP-conjugated goat anti-rabbit IgG (Bio-Rad), HRP-conjugated donkey anti-goat IgG (Santa Cruz), HRP-conjugated donkey anti-rat IgG (Abcam).

### Yeast two-hybrid assays

Yeast two-hybrid vectors were transformed separately into haploid strains SKY661 and SKY662 and selected on appropriate synthetic dropout medium. Strains were mated and streaked for single diploid colonies on medium lacking tryptophan and leucine. Single colonies were grown overnight in selective medium containing 2% glucose. Cultures were diluted in fresh medium containing 2% galactose and 1% raffinose and grown until log phase (4 h). Cells were lysed and quantitative β-galactosidase assay was performed using ONPG substrate following standard protocols (Clontech Laboratories). For two-hybrid experiments in meiotic conditions, after overnight culture in selective medium with glucose, cultures were washed twice then incubated with 2% KOAc for 20 hours to induce meiosis.

### Spo11-oligo sequencing and S1-seq

Processing of cell lysates for Spo11 oligo purification and sequencing library preparation were essentially as described (Lam et al., 2017). Paired-end Illumina sequencing was performed at the Integrated Genomics Operation facility at MSKCC. Clipping of library adapters and mapping of reads was performed by the Bioinformatics Core Facility (MSKCC) using a custom pipeline as described (Pan et al., 2011). Statistical analyses were performed using R (http://www.r-project.org/). Sequence read totals and mapping statistics are described in **Supplementary Table 5**. After mapping, the reads were separated into unique and multiple mapping sets, but only uniquely mapping reads were analyzed. Each map was normalized to the total number of uniquely mapped reads (reads per million, RPM; excluding reads mapping to mitochondrial DNA or the 2µ plasmid), and then maps for biological replicates were averaged. In analyses evaluating the fold change in DSB activity, the ratio was scaled according to the decrease in DSBs as measured through quantification of Spo11-oligo complexes (0.4). Sequence reads and compiled Spo11-oligo maps have been submitted to the Gene Expression Omnibus (GEO) repository (accession number pending).

For S1-seq, yeast cultures in the *sae2* background were harvested 4 hours after meiotic induction, washed with 50 mM EDTA pH 8.0 and stored at -80°C. DNA preparation in plugs and generation of sequencing libraries were done as described (Mimitou and Keeney, 2018). Signal mapping to the rDNA locus was masked. In addition, to reduce background signal, a thresholding was applied, calculated from the frequency of Spo11-oligos in coldest 10% of coding regions.

## End Matter

### Author Contributions and Notes

C.C.B., S.E.T. and S.K. designed research, C.C.B., S.E.T., S.P., E.P.M. and E.A-P. performed research, all authors analyzed data; C.C.B and S.K. wrote the paper, with input from all authors. The authors declare no conflict of interest.

## Acknowledgments

We thank Matthew Brendel from the Molecular Cytology core facility at MSKCC for performing the AFM experiments. We thank Ronald Hendrickson and Elizabeth Chang from the Microchemistry and Proteomics core facility at MSKCC for assistance with the XL-MS experiments. MSKCC core facilities are supported by NCI Cancer Center support grant P30 CA08748. We thank Ewa Folta-Stogniew from the Biophysics Resource of Keck Facility at Yale University for the SEC-MALS experiments. The SEC-LS/UV/RI instrumentation was supported by NIH Award Number 1S10RR023748-01. EPM was supported in part by a Helen Hay Whitney Foundation fellowship. Work in the SK lab was supported principally by the Howard Hughes Medical Institute and in part by NIH grant R35 GM118092. Work in the JMB lab was funded by NCI grant R01-CA0777373.

## Supplementary files

**Supplementary Figure 1:**
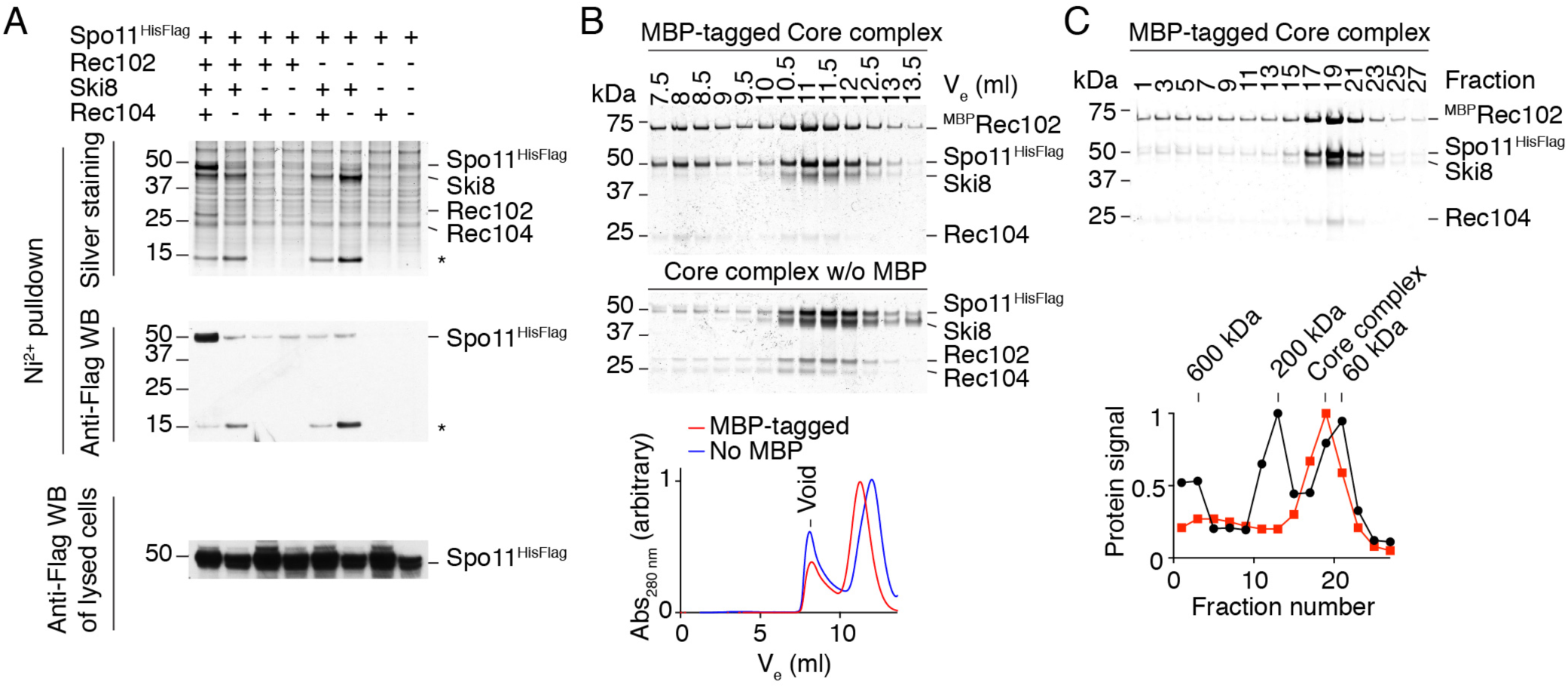
Co-expression of core complex subunits, size exclusion chromatography and glycerol gradient sedimentation analyses of the core complex. A. Silver-stained SDS-PAGE gels (top) and anti-Flag western blot (WB; center) of Spo11 complexes after purification on nickel resin. Absence of Rec102, Rec104, or Ski8 leads to poor solubility of Spo11. Bottom: anti-Flag western blot of lysed Sf9 cells showing Spo11 expression levels. Asterisks: C-terminal truncation of Spo11 that retains the affinity tag and interaction with Ski8. B. Size exclusion chromatography of purified core complex with and without MBP tag on Rec102. Silver-stained SDS-PAGE gels of eluted fractions are shown above, with chromatograms from absorption at 280 nm below. C. Glycerol gradient sedimentation of MBP-tagged Spo11 core complexes. The silver-stained SDS-PAGE gel shows fractions collected from the bottom of the gradient. Quantification of protein signal from two independent experiments is shown together with molecular weight markers run on a separate gradient and quantified by Bradford assay. Note: Material in the void volume (panel B) and at the bottom of the glycerol gradient (panel C) lacks Ski8, which is consistent with Ski8 being required for solubility.

**Supplementary Figure 2:**
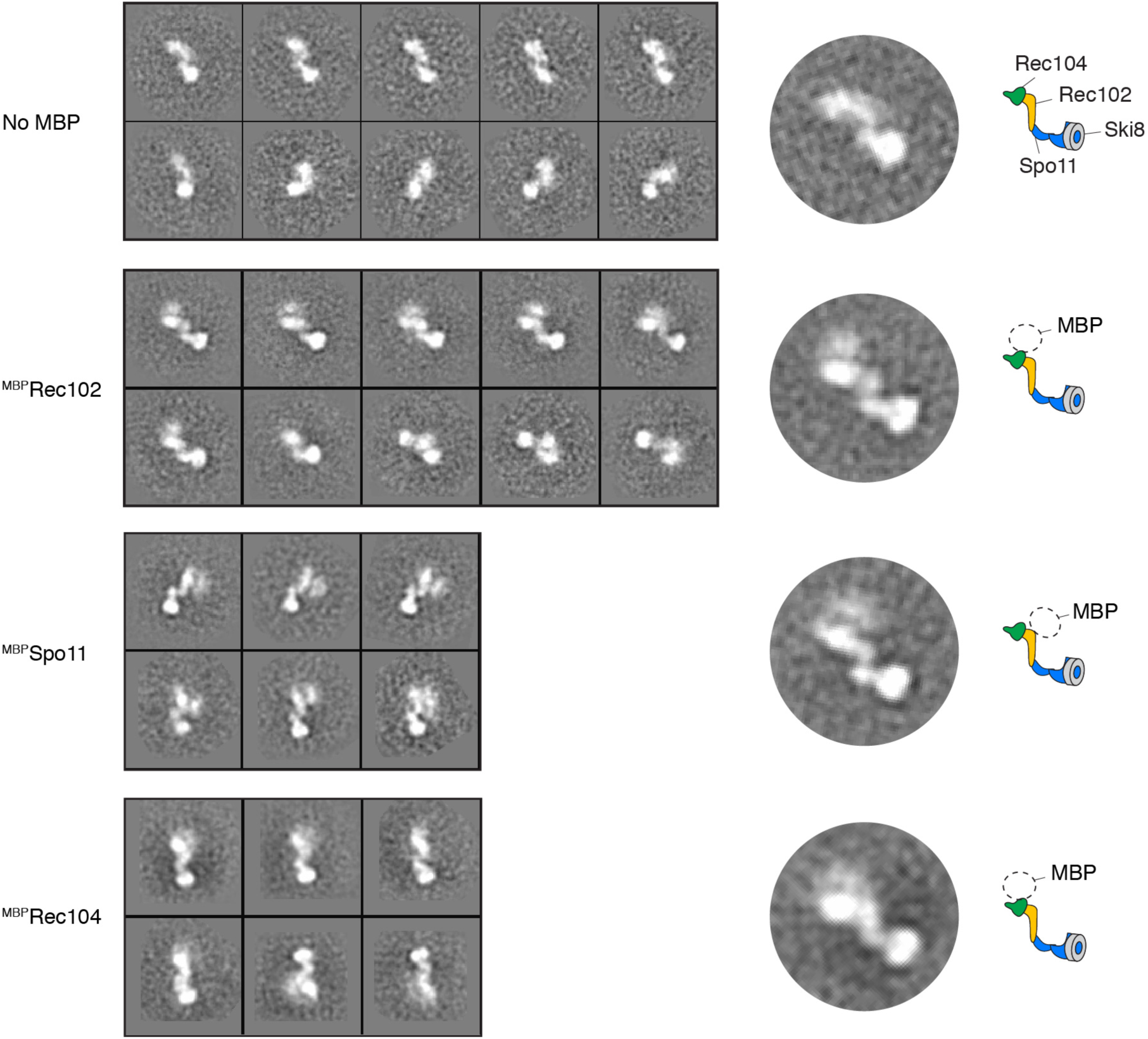
2D class averages of nsEM images with different versions of the core complex. Core complexes without MBP or with MBP fused at the N-terminus of Rec102, Spo11 or Rec104 are shown. A cartoon of the presumed arrangement of the subunits and the position of the MBP electron density is shown. With the MBP-tagged Spo11 construct, the electron density of MBP is located at a similar position to the Rec102- or Rec104-tagged constructs. This is consistent with the observation that the N-terminus of Spo11 frequently crosslinks with Rec104 (pink lines in **Figure 2A**), suggesting that the N-terminus of Spo11, absent from the structural model, is flexible and perhaps directly contacts Rec102/Rec104. Complexes with MBP-tagged Ski8 were not well behaved and could not be purified.

**Supplementary Figure 3:**
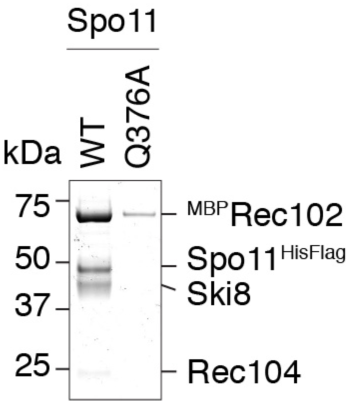
The interaction between Ski8 and Spo11 is important for the integrity of the complex. SDS-PAGE analysis of core complexes purified with wild-type Spo11 or the Ski8-interaction deficient Q376A mutant. Equivalent percentages of the total protein purified from similar amounts of Sf9 extract were loaded in each lane, demonstrating the poor yield when the Spo11–Ski8 interaction is compromised.

**Supplementary Figure 4:**
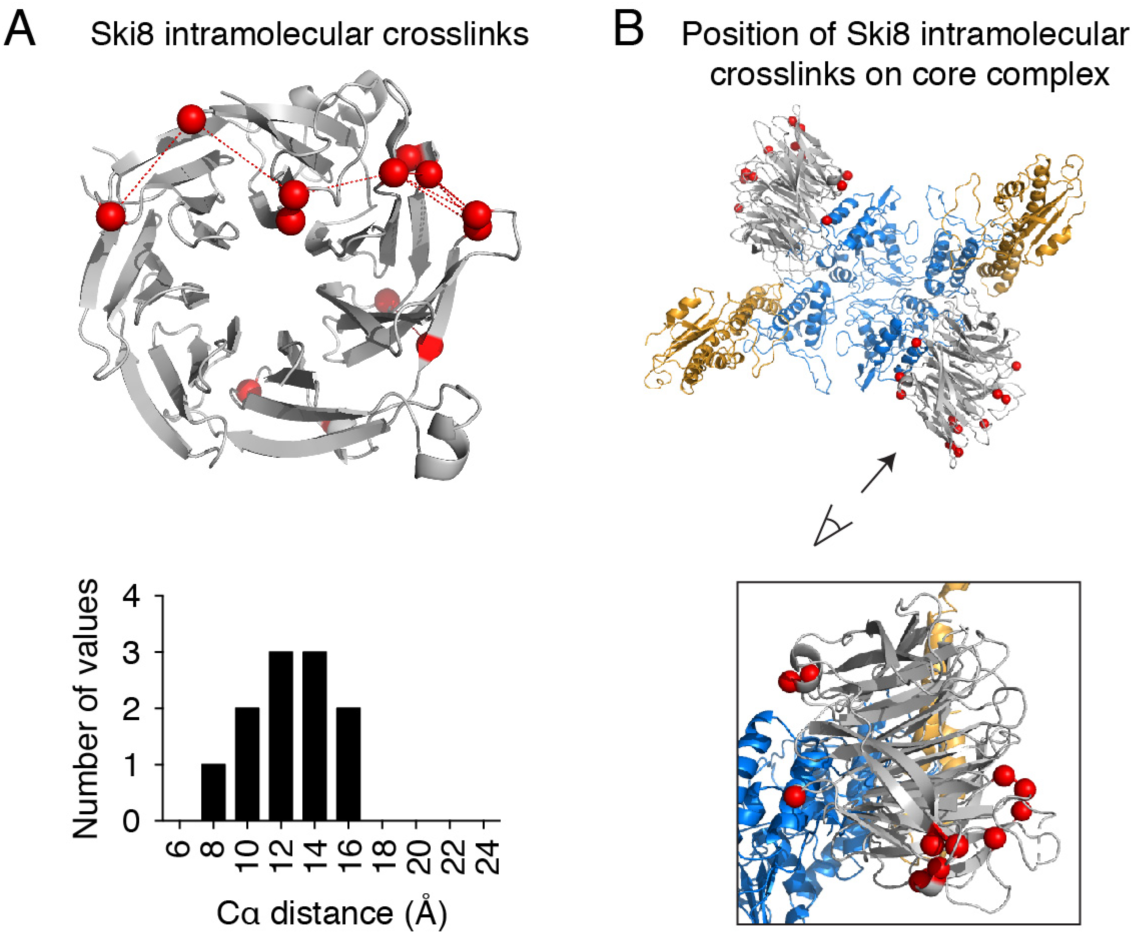
Intramolecular crosslinks within Ski8 validate the XL-MS results. A. Ski8 intramolecular crosslinks modeled on the structure of Ski8. The histogram shows the frequency of XL-MS events as a function of distance between the α-carbons (Cα) of the crosslinked lysines (red spheres). The crosslinkable limit of DSS is 27.4 Å. B. Ski8 intramolecular crosslinks modeled on the core complex show that the crosslinked residues are away from the interaction surface with Spo11.

**Supplementary Figure 5:**
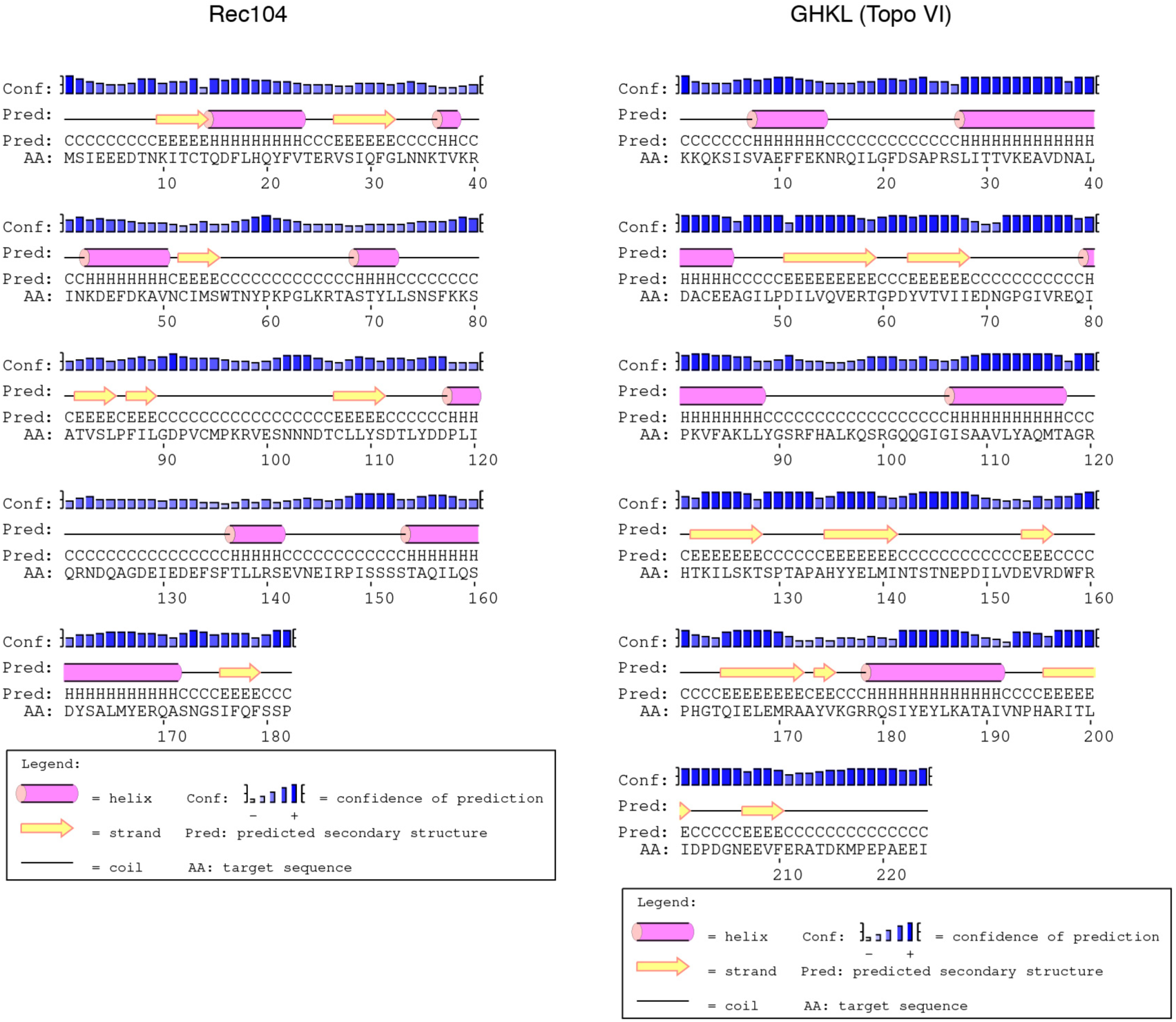
Secondary structure prediction of Rec104. Secondary structure predictions for Rec104 and the GHKL domain of Topo VIB were generated by PsiPred (McGuffin et al., 2000).

**Supplementary Figure 6:**
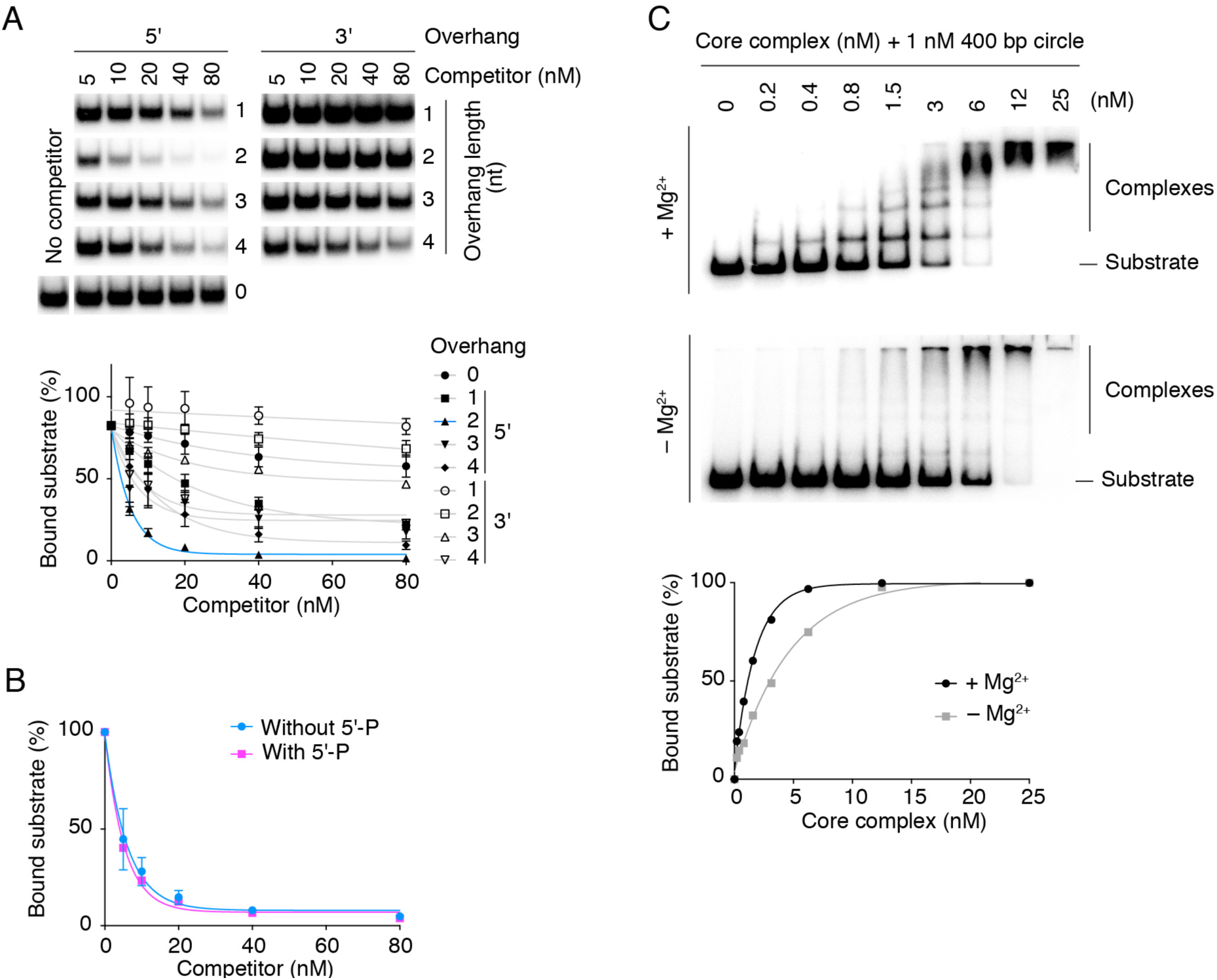
DNA-binding properties of the core complex. A. Competition experiment using a labeled 25-bp hairpin substrate with 5′-TA overhang in the presence of unlabeled substrates with various overhang configurations. EMSA gel bands of bound labeled substrate are shown. Mean and ranges from two experiments are plotted. The substrate with a 2-nucleotide 5′ overhang is the most effective competitor. B. Competition experiment using a labeled 25-bp hairpin substrate with 5′-TA overhang in the presence of unlabeled competitor substrates with or without 5’ phosphate. Error bars represent ranges from two experiments. C. EMSA of core complex binding to 400-bp mini-circles in the presence or absence of Mg^2+^. For the top panel, binding reactions contained 5 mM Mg^2+^ and the gel and electrophoresis buffer contained 0.5 mM Mg^2+^. For the bottom panel, the binding reactions, gel, and buffer contained 1 mM EDTA.

**Supplementary Figure 7:**
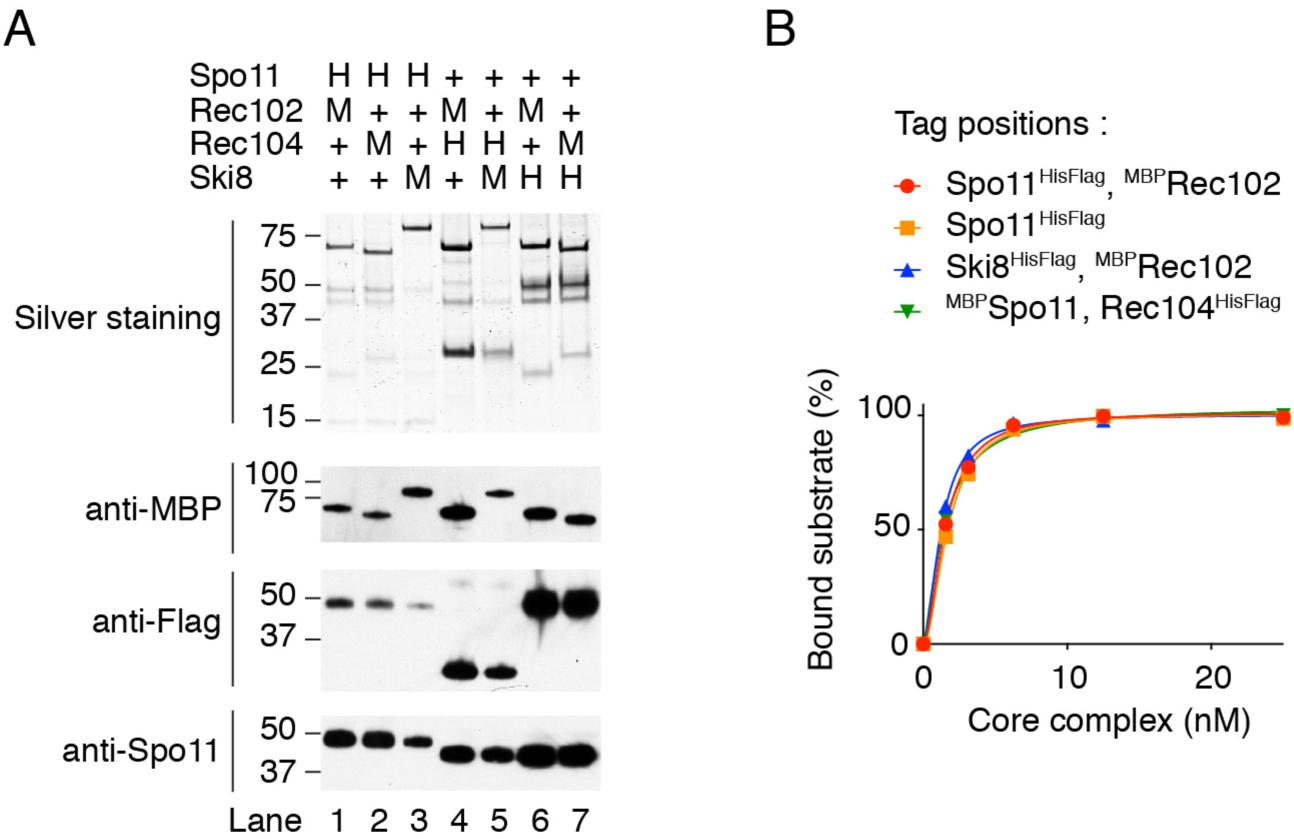
Affinity purification of different combinations of tagged complexes and comparison of DNA-binding activities. A. Purification of core complexes that carry combinations of HisFlag (H) and MBP (M) tags on different subunits. All combinations yielded soluble Spo11 (western blot, bottom panel). B. Comparison of the DNA-binding activity of core complexes that carried affinity tags on different subunits. All tagged complexes assayed had similar DNA-binding activities.

**Supplementary Figure 8:**
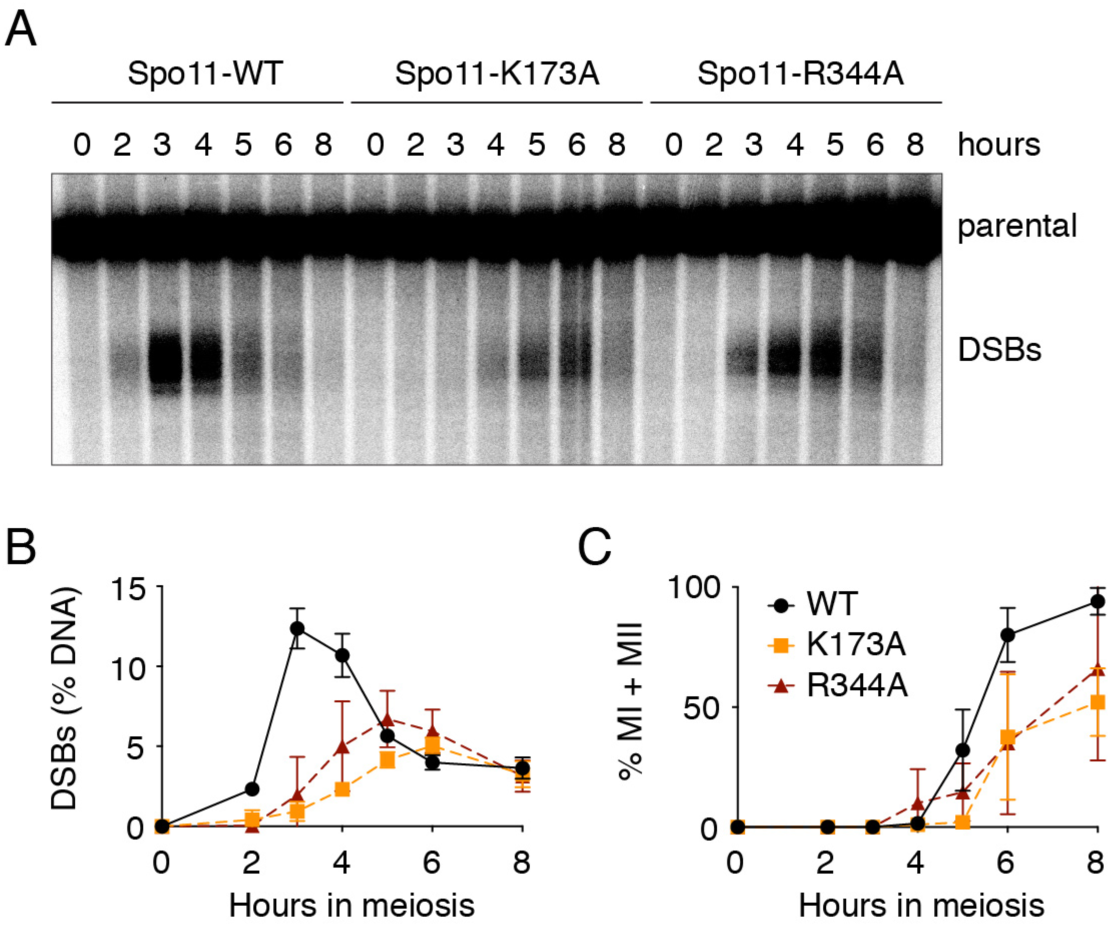
*In vivo* analyses of Spo11 DNA-binding mutants. A. Southern blot analysis of meiotic DSB formation at the *CCT6* hotspot in strains expressing wild-type (WT) Spo11 or the K173A or R344A mutant proteins. B. Quantification of DSB formation at the *CCT6* hotspot. Error bars represent the range from two experiments. C. Meiotic progression. MI + MII indicates the fraction of cells that have undergone the first or both meiotic divisions, as scored by DAPI staining.

**Supplementary Figure 9:**
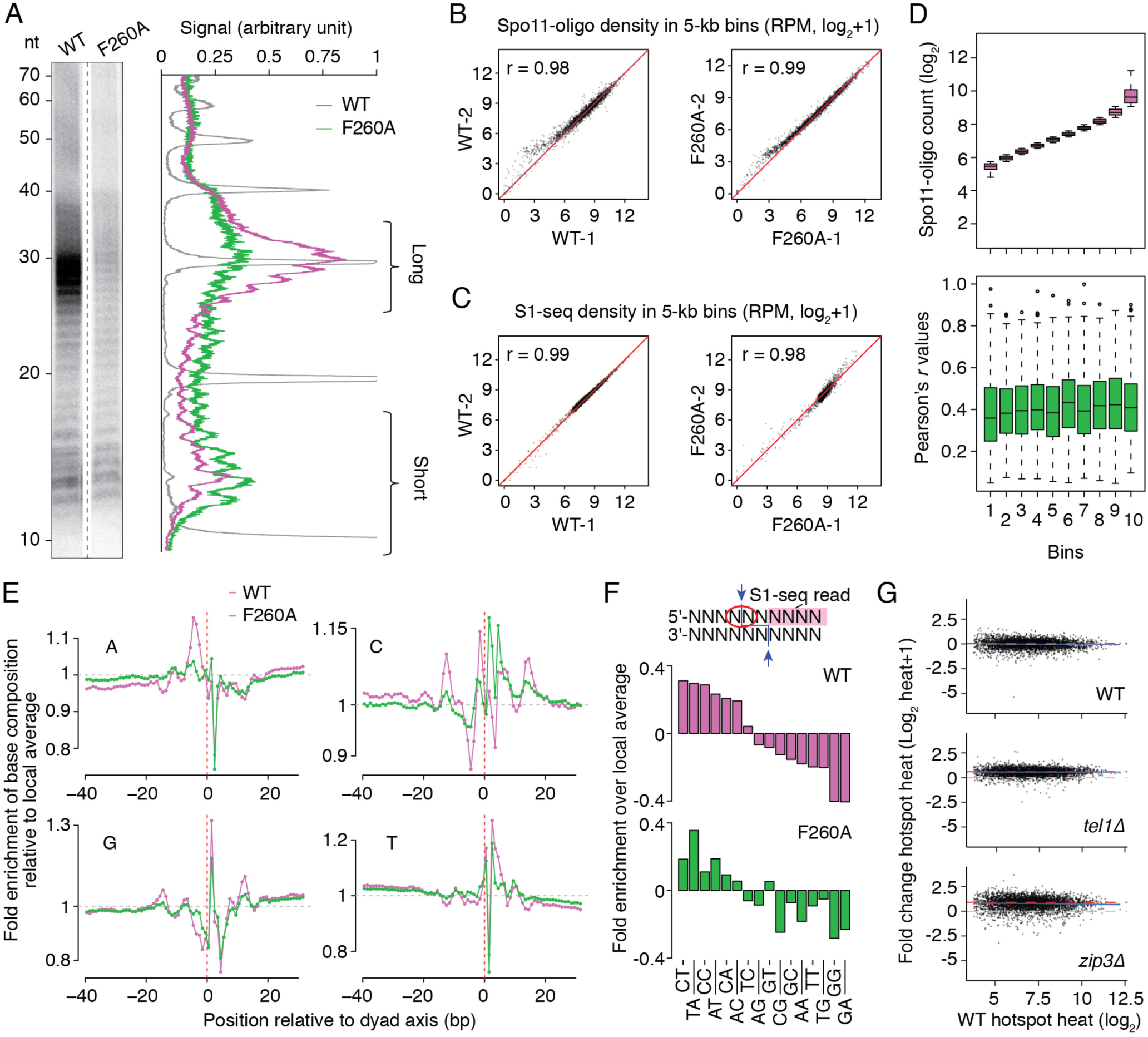
Genome-wide analyses of DSB formation in the F260A mutant. A. Relative enrichment of the short Spo11-oligo class in F260A. Deproteinized, labeled oligos were separated by denaturing gel electrophoresis. Lane profiles are shown on the right. A 10-nt ladder is plotted in grey. B, C. Reproducibility of DSB maps. Correlations of Spo11-oligo counts (B) and S1-seq counts (C) within hotspots between two biological replicates of Spo11 wild type and F260A are plotted. Pearson’s *r* between datasets is indicated. D. Changed DSB distribution in F260A is not correlated with hotspot strength. Spo11 hotspots were binned according to oligo counts in wild type. Boxplots show the distribution of Pearson’s *r* values comparing within-hotspot Spo11-oligo distributions between wild type and F260A, as in **Figure 8E**. The thick horizontal bars are medians, box edges are upper and lower quartiles, whiskers indicate values within 1.5 fold of interquartile range, and points are outliers. E. Base composition in S1-seq maps. The big spike in the G map at +2 is partially because this is the complement of the preferred C 5′ of the scissile phosphate, but it is also the first base of the ligation junction (and end-most base after S1 digestion), so the degree to which there is enrichment to the right but not left of the dyad axis probably reflects a modest end-bias in library prep in S1-seq. F. Spo11 preference at the scissile phosphate (dinucleotide indicated by the red circle). G. Changes in absolute hotspot strength as a function of hotspot strength in wild type (compare with **Figure 8J**). Log-fold changes are for comparison of a consensus wild-type map to wild-type datasets generated in this study or to *tel1Δ* and *zip3Δ* mutants. The *tel1Δ* and *zip3Δ* data were scaled by factors of 1.5 and 1.8, respectively based on measured increases in total Spo11-oligo complexes (Thacker et al., 2014; Mohibullah and Keeney, 2017)

**Supplementary Figure 10:**
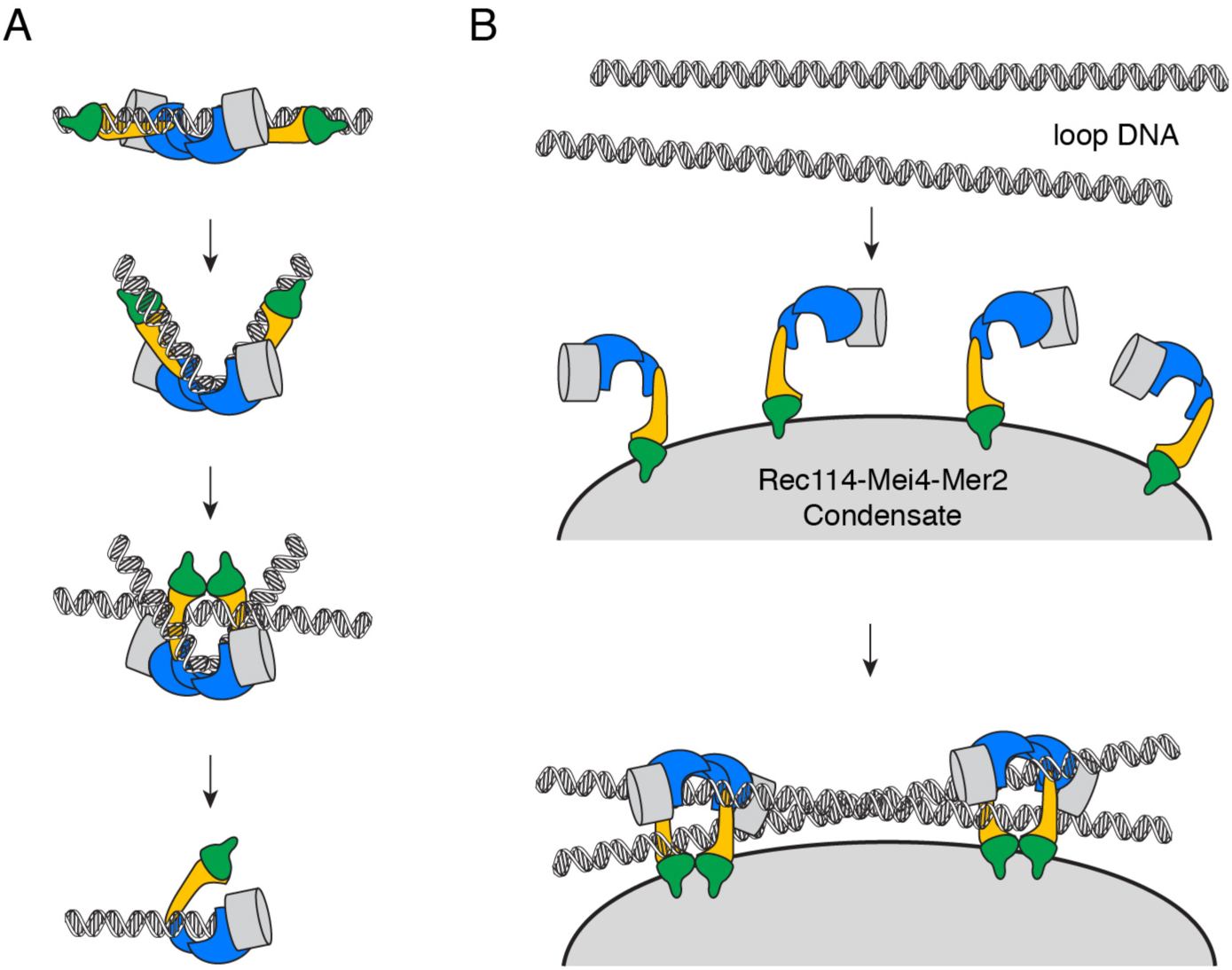
Model of Spo11-induced break formation. A. AFM experiments suggest a model where the core complex binds a DNA duplex, bends it, then traps a second duplex. Perhaps DNA cleavage happens in the context of a trapped DNA junction, similar to Topo VI. After cleavage, Spo11 remains covalently attached to the DNA end through covalent and non-covalent interactions. B. Model of assembly of the DSB machinery. DNA-driven condensation by Rec114–Mei4–Mer2 is proposed to provide a platform that recruits the core complex, where it engages its DNA substrate (Claeys Bouuaert et al., 2020).

Supplementary Tables 1: XL-MS data.

**Supplementary Table 2:**
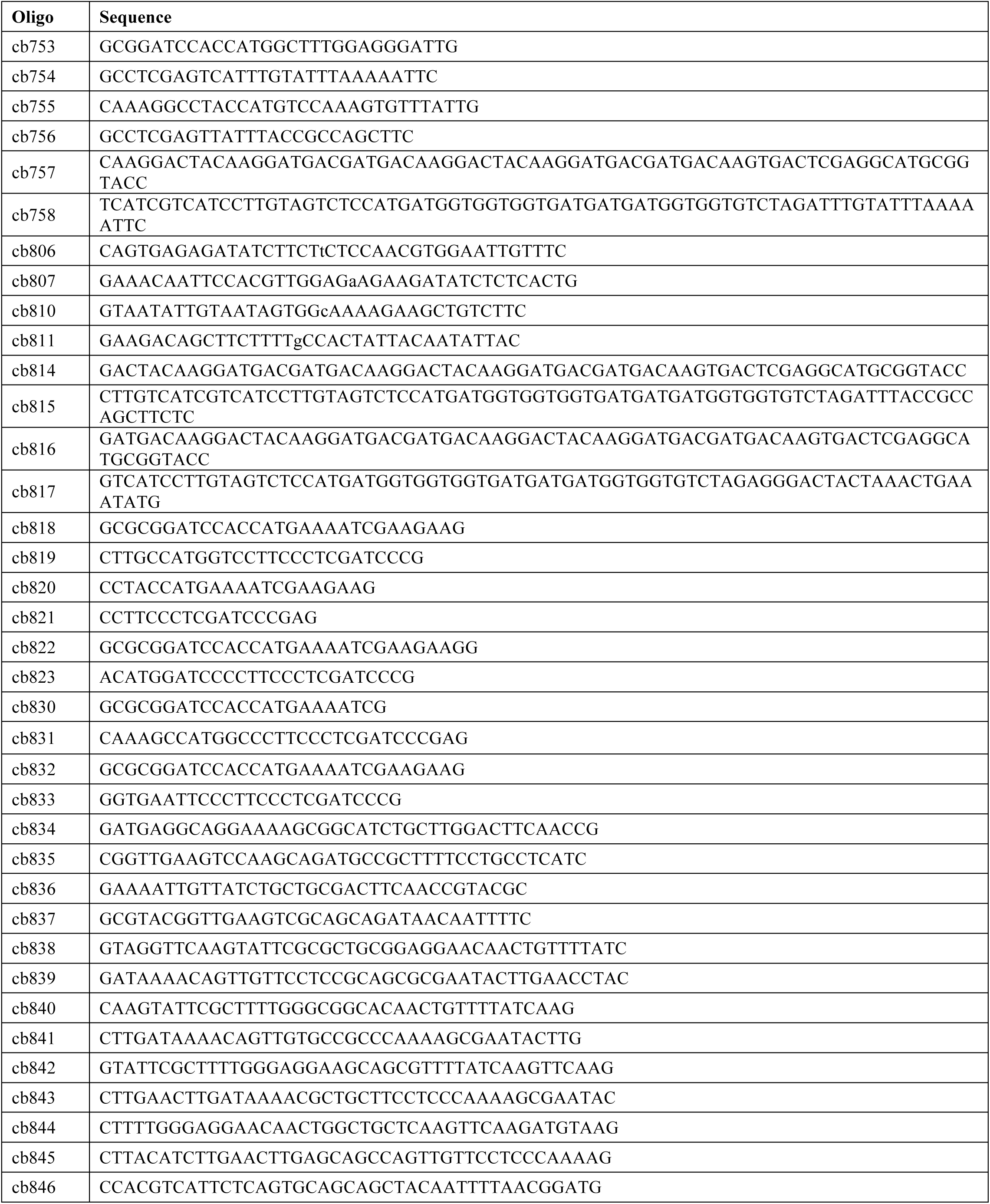
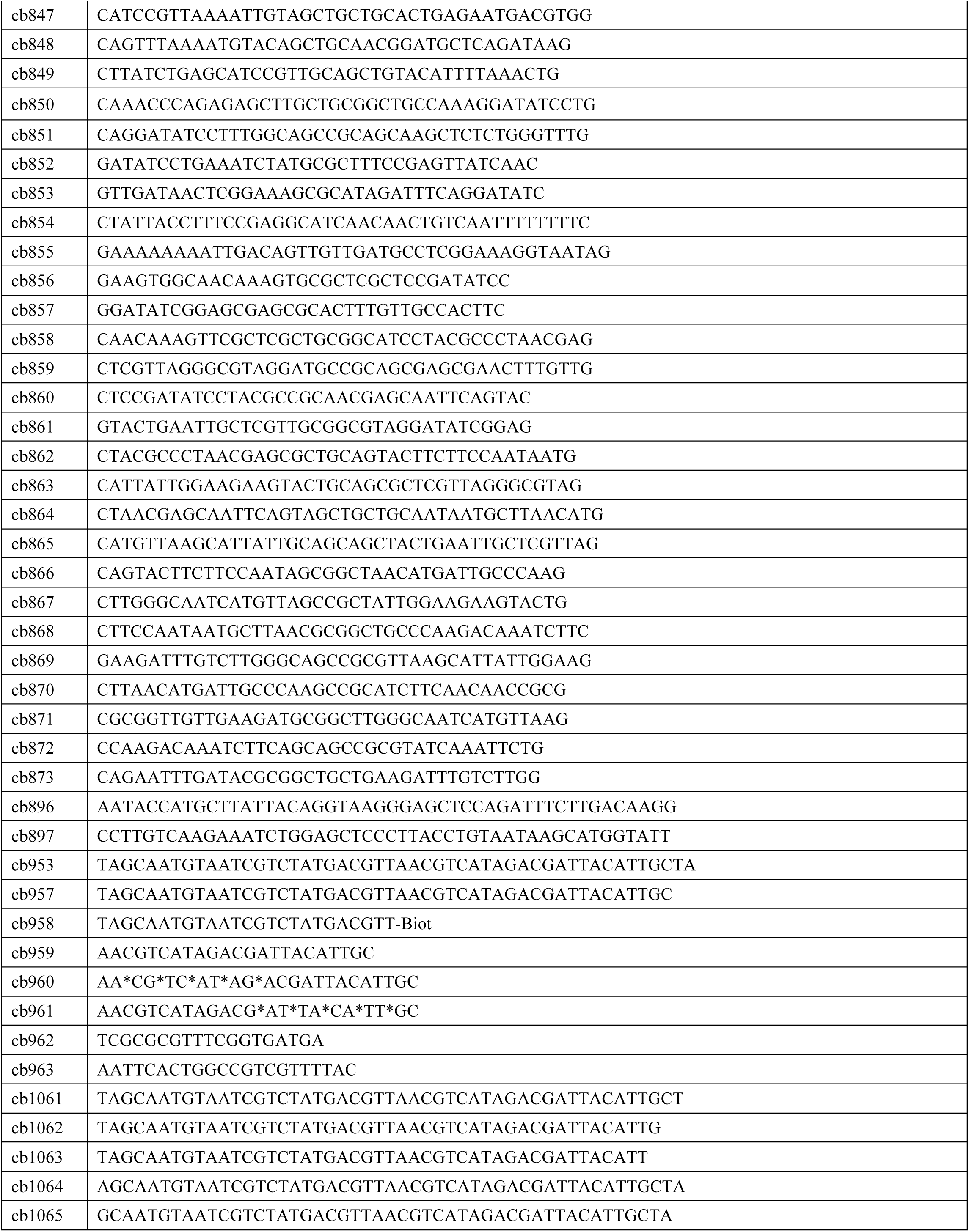
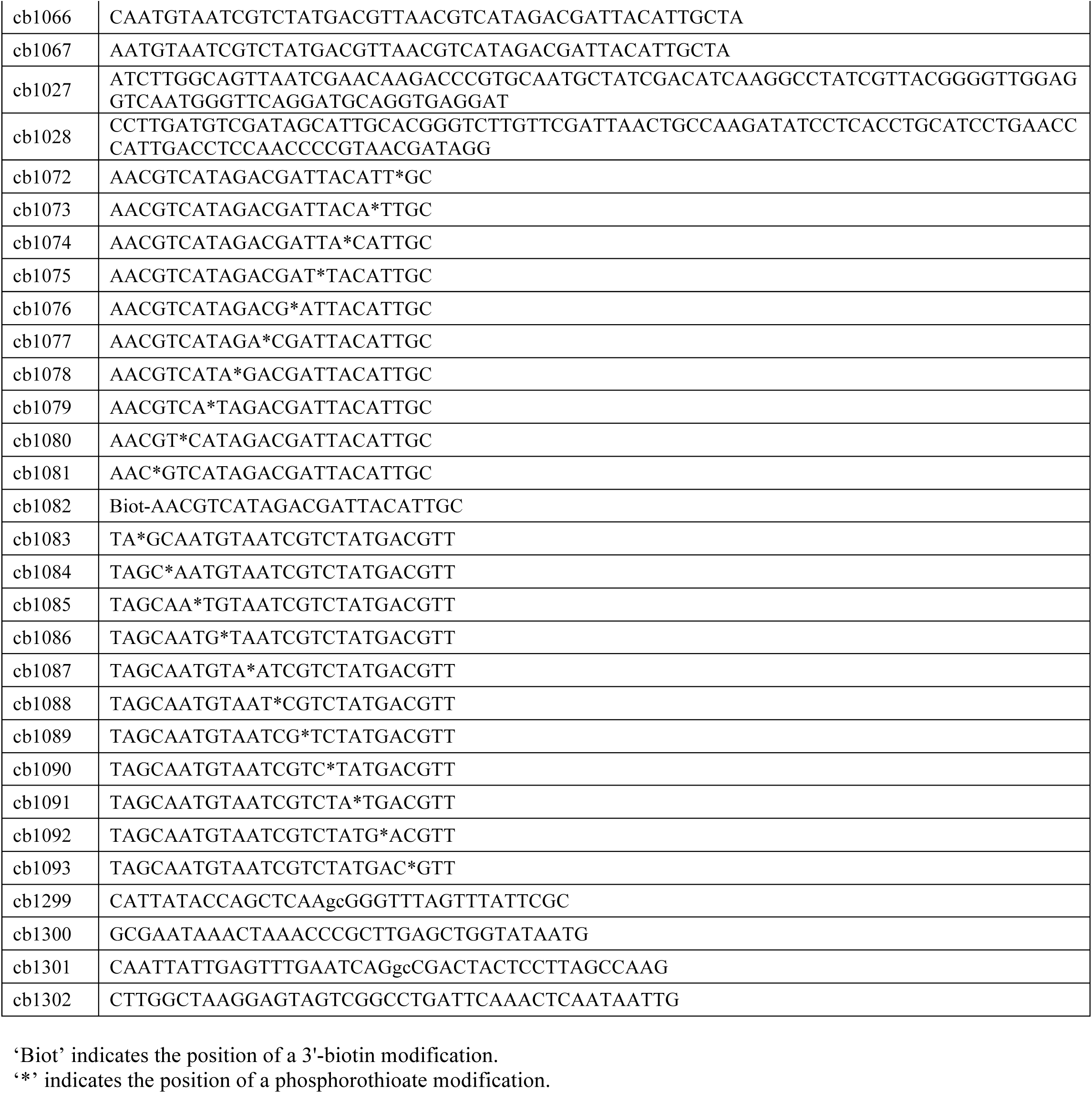
Oligonucleotides used in this study.

**Supplementary Table 3:**
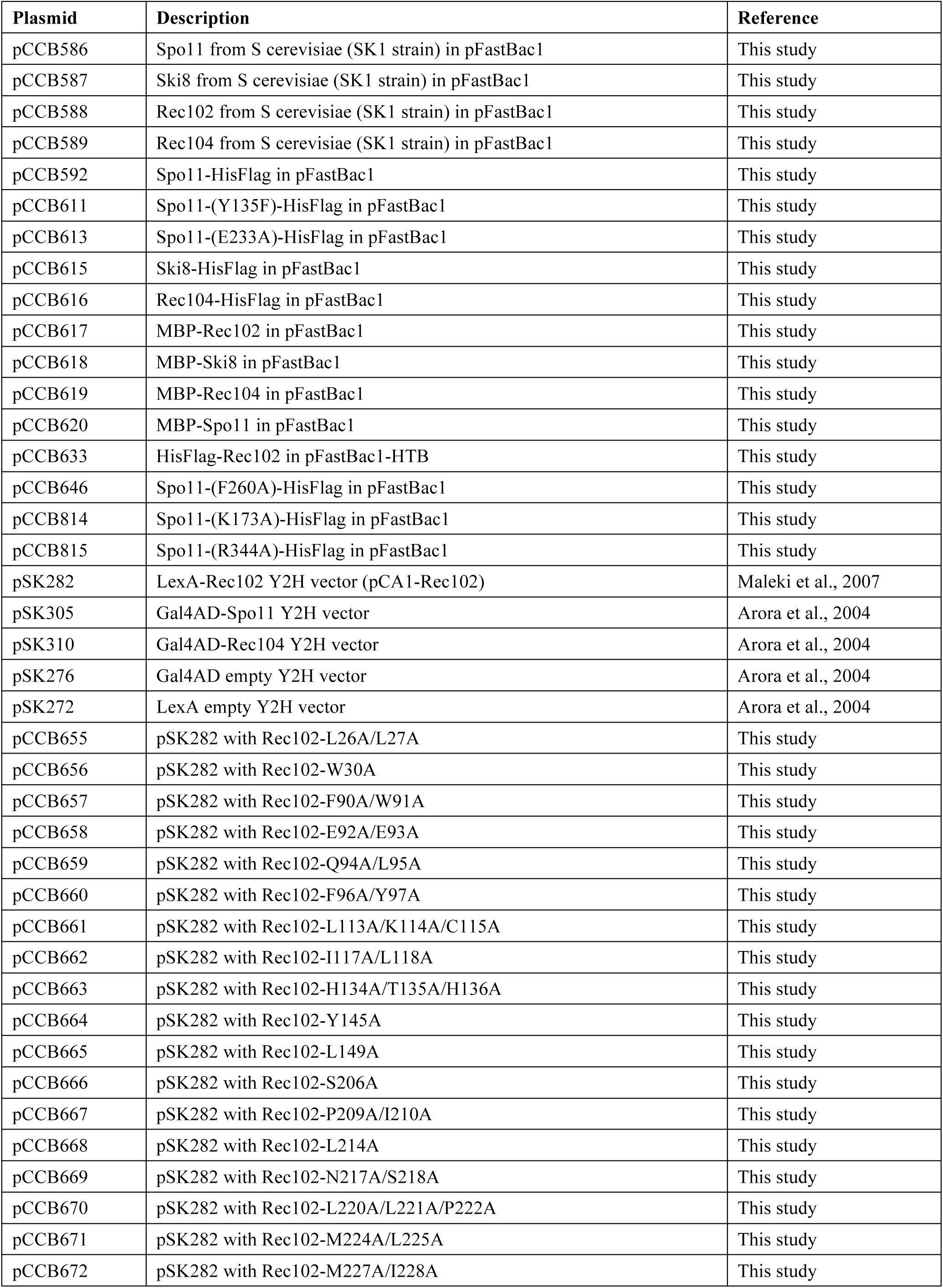
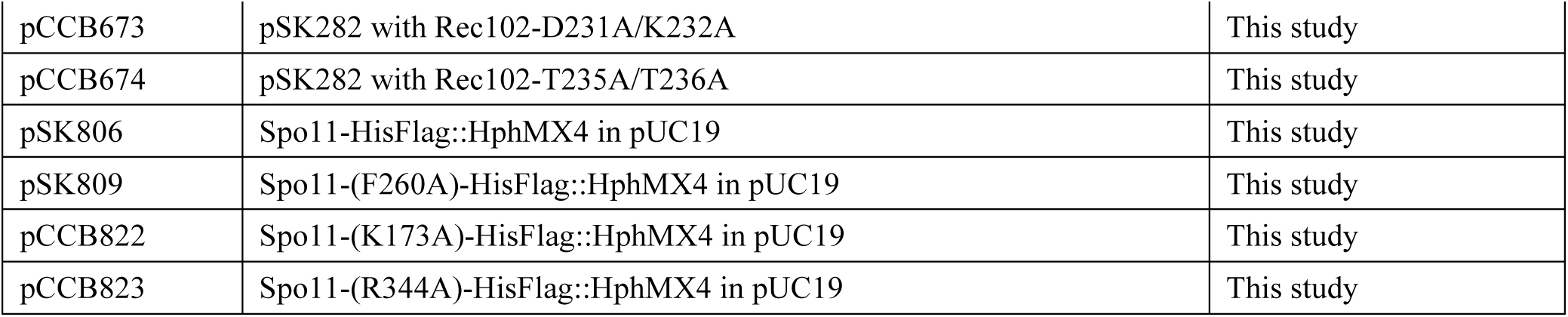
Plasmids used in this study.

| **Plasmid** | **Description** | **Reference** |
| --- | --- | --- |
| pCCB586 | Spo11 from *S cerevisiae* (SK1 strain) in pFastBac1 | This study |
| pCCB587 | Ski8 from *S cerevisiae* (SK1 strain) in pFastBac1 | This study |
| pCCB588 | Rec102 from *S cerevisiae* (SK1 strain) in pFastBac1 | This study |
| pCCB589 | Rec104 from *S cerevisiae* (SK1 strain) in pFastBac1 | This study |
| pCCB592 | Spo11-HisFlag in pFastBac1 | This study |
| pCCB611 | Spo11-(Y135F)-HisFlag in pFastBac1 | This study |
| pCCB613 | Spo11-(E233A)-HisFlag in pFastBac1 | This study |
| pCCB615 | Ski8-HisFlag in pFastBac1 | This study |
| pCCB616 | Rec104-HisFlag in pFastBac1 | This study |
| pCCB617 | MBP-Rec102 in pFastBac1 | This study |
| pCCB618 | MBP-Ski8 in pFastBac1 | This study |
| pCCB619 | MBP-Rec104 in pFastBac1 | This study |
| pCCB620 | MBP-Spo11 in pFastBac1 | This study |
| pCCB633 | HisFlag-Rec102 in pFastBac1-HTB | This study |
| pCCB646 | Spo11-(F260A)-HisFlag in pFastBac1 | This study |
| pCCB814 | Spo11-(K173A)-HisFlag in pFastBac1 | This study |
| pCCB815 | Spo11-(R344A)-HisFlag in pFastBac1 | This study |
| pSK282 | LexA-Rec102 Y2H vector (pCA1-Rec102) | Maleki et al., 2007 |
| pSK305 | Gal4AD-Spo11 Y2H vector | Arora et al., 2004 |
| pSK310 | Gal4AD-Rec104 Y2H vector | Arora et al., 2004 |
| pSK276 | Gal4AD empty Y2H vector | Arora et al., 2004 |
| pSK272 | LexA empty Y2H vector | Arora et al., 2004 |
| pCCB655 | pSK282 with Rec102-L26A/L27A | This study |
| pCCB656 | pSK282 with Rec102-W30A | This study |
| pCCB657 | pSK282 with Rec102-F90A/W91A | This study |
| pCCB658 | pSK282 with Rec102-E92A/E93A | This study |
| pCCB659 | pSK282 with Rec102-Q94A/L95A | This study |
| pCCB660 | pSK282 with Rec102-F96A/Y97A | This study |
| pCCB661 | pSK282 with Rec102-L113A/K114A/C115A | This study |
| pCCB662 | pSK282 with Rec102-I117A/L118A | This study |
| pCCB663 | pSK282 with Rec102-H134A/T135A/H136A | This study |
| pCCB664 | pSK282 with Rec102-Y145A | This study |
| pCCB665 | pSK282 with Rec102-L149A | This study |
| pCCB666 | pSK282 with Rec102-S206A | This study |
| pCCB667 | pSK282 with Rec102-P209A/I210A | This study |
| pCCB668 | pSK282 with Rec102-L214A | This study |
| pCCB669 | pSK282 with Rec102-N217A/S218A | This study |
| pCCB670 | pSK282 with Rec102-L220A/L221A/P222A | This study |
| pCCB671 | pSK282 with Rec102-M224A/L225A | This study |
| pCCB672 | pSK282 with Rec102-M227A/I228A | This study |
| pCCB673 | pSK282 with Rec102-D231A/K232A | This study |
| pCCB674 | pSK282 with Rec102-T235A/T236A | This study |
| pSK806 | Spo11-HisFlag::HphMX4 in pUC19 | This study |
| pSK809 | Spo11-(F260A)-HisFlag::HphMX4 in pUC19 | This study |
| pCCB822 | Spo11-(K173A)-HisFlag::HphMX4 in pUC19 | This study |
| pCCB823 | Spo11-(R344A)-HisFlag::HphMX4 in pUC19 | This study |

**Supplementary Table 4:** Yeast strains used in this study.

| Strain | Genotype |
| --- | --- |
| SKY661 | *MATa, ho::LYS2, lys2, leu2::hisG, trp1::hisG, ndt80::kanMX, LexA(op)-LacZ::URA3* |
| SKY662 | *MATα, ho::LYS2, lys2, leu2::hisG, trp1::hisG, ndt80::kanMX, LexA(op)-LacZ::URA3* |
| SKY5788 | *MATa, ho::LYS2, lys2, ura3, leu2::hisG, arg4-Nsp, nuc1Δ::LEU2, SPO11-His6-flag3-HphMX4* |
| SKY5789 | *MATα, ho::LYS2, lys2, ura3, leu2::hisG, arg4-Nsp, nuc1Δ::LEU2, SPO11-His6-flag3-HphMX4* |
| SKY5797 | *MATa, ho::LYS2, lys2, ura3, leu2::hisG, arg4-Nsp, nuc1Δ::LEU2, SPO11-(F260A)-His6-flag3-HphMX4* |
| SKY5798 | *MATα, ho::LYS2, lys2, ura3, leu2::hisG, arg4-Nsp, nuc1Δ::LEU2, SPO11-(F260A)-His6-flag3-HphMX4* |
| SKY5806 | *MATa, ho::LYS2, lys2, ura3, leu2::hisG, arg4-Nsp, nuc1Δ::LEU2, SPO11-His6-flag3-HphMX4, sae2Δ::KanMX6* |
| SKY5807 | *MATα, ho::LYS2, lys2, ura3, leu2::hisG, arg4-Nsp, nuc1Δ::LEU2, SPO11-His6-flag3-HphMX4, sae2Δ::KanMX6* |
| SKY6621 | *MATa, ho::LYS2, lys2, ura3, leu2::hisG, arg4-Nsp, nuc1Δ::LEU2, SPO11-(F260A)-His6-flag3-HphMX4, sae2::KanMX6* |
| SKY6622 | *MATα, ho::LYS2, lys2, ura3, leu2::hisG, arg4-Nsp, nuc1Δ::LEU2, SPO11-(F260A)-His6-flag3-HphMX4, sae2::KanMX6* |
| SKY6615 | *MATa, ho::LYS2, lys2, ura3, leu2::hisG, SPO11(K173A)-His6-flag3-HphMX4* |
| SKY6616 | *MATα, ho::LYS2, lys2, ura3, leu2::hisG, SPO11(K173A)-His6-flag3-HphMX4* |
| SKY6617 | *MATa, ho::LYS2, lys2, ura3, leu2::hisG, SPO11(R344A)-His6-flag3-HphMX4* |
| SKY6618 | *MATα, ho::LYS2, lys2, ura3, leu2::hisG, SPO11(R344A)-His6-flag3-HphMX4* |

All strains are from the SK1 background.

